# Simultaneous Control of Infection and Inflammation by Keratin-Derived Antibacterial Peptides (KAMPs) Targeting TLRs and Co-Receptors

**DOI:** 10.1101/2021.01.18.427180

**Authors:** Yan Sun, Jonathan Chan, Karthikeyan Bose, Connie Tam

## Abstract

The use and the timing of initiation of steroids for controlling unwanted infectious inflammation are major clinical dilemmas due to their possible adverse effects including delayed microbial clearance and wound healing. Compounding this difficulty is the continued emergence of drug-resistant bacteria; yet anti-infective strategies aiming at augmenting inflammatory responses to facilitate bacterial killing cannot be used to treat infections involving vulnerable tissues. As is the case with bacterial keratitis, excessive inflammation jeopardizes corneal transparency leading to devastating vision loss. Hence, a two-pronged remedy possessing both anti-infective and anti-inflammatory properties would be helpful for tackling antibiotic resistance and enabling prompt inflammation control at once. Using murine primary neutrophils, macrophages and sterile corneal inflammation models, we found that non-toxic and pro-healing human keratin 6a-derived antimicrobial peptides (KAMPs) with a native 10-or 18-amino-acid sequence suppress LTA- and LPS-induced NF-кB and IRF3 activation, proinflammatory cytokine production, as well as phagocyte recruitment, independently of their bactericidal function. Mechanistically, direct binding of KAMPs to cell surface TLR2 and TLR co-receptors CD14 and MD-2 not only blocks their bacterial ligand docking sites, but also reduces cell surface availability of TLR2 and TLR4 through promotion of receptor endocytosis. Benefitting from the dual functions of topical KAMPs, experimental bacterial keratitis caused was effectively prevented or controlled, as evidenced by significant reductions of corneal opacification and inflammatory cell infiltration in addition to enhanced bacterial clearance. These findings reveal multiple TLR-targeting activities of KAMPs and demonstrate their therapeutic potential as a multifunctional drug for managing sterile and infectious inflammatory diseases.

**One Sentence Summary:** Bifunctional native keratin peptides allow concurrent alleviation of inflammation and infection to avoid functional damages in vulnerable tissues.

## INTRODUCTION

Although infection-driven immune cell infiltration is a natural inflammatory response to fight invading microbes, it can easily cause serious damage to vulnerable tissues leading to permanent organ dysfunction. To reduce the risk of long-term disability and mortality, the infection itself and associated acute inflammation should both be controlled as early as possible *(1-3)*. Unfortunately, this treatment goal is hampered not only by the ever-growing problem of antibiotic resistance, but also by a lack of safer anti-inflammatory drugs. Precisely, even though corticosteroids are potent immunosuppressive agents that alleviate inflammation, their usage in conjunction with active antimicrobial therapy has long remained an important clinical dilemma due to their potential harmful effects including impairment of microbial clearance and severe complications. Indeed, the use of adjunctive corticosteroids in infections is far from optimal as it can be beneficial or harmful depending on a number of conditions, such as comorbidity, dosing, timing of treatment initiation, and type of causative pathogen *(4-7)*. Undoubtedly, advance new therapeutic options are needed for simultaneous control of infection and inflammation.

Bacterial keratitis is among the infectious inflammatory diseases that affect tissues functionally vulnerable to the deleterious effects of inflammation. In this instance, scarring of the cornea impacts its optical clarity and proper refractive shape that are required for good vision, leading to devastating visual impairment or blindness. The first-line treatment is topical antibiotics; particularly fortified vancomycin and fluoroquinolone monotherapy are the most common empirical choices *(7, 8)*. However, both are limited by their respective drawbacks of high ocular toxicity and low susceptibility of methicillin-resistant *Staphylococcus aureus (8, 9)*. Notably, rising antibiotic resistance among ocular pathogens is significantly associated with worse clinical outcomes *(10, 11)*. Even if infectious organisms are susceptible to treatment, inflammation can continue to progress causing severe tissue injury and opacification that irreversibly impair vision *(12)*. To mitigate inflammation-associated damages, current guidelines recommend initiation of adjunctive topical corticosteroids for culture-positive non-Nocardia bacterial keratitis 24 to 48 hours after initial antibiotic administration when infection is under control *(13)*, albeit their potential negative effect on re-epithelialization of the cornea *(14-16)*. Notably, early addition of corticosteroids to the treatment course is necessary for improved visual outcomes *(3)*, as in the case of bacterial meningitis, hearing loss and neurologic damage can be prevented only when adjunctive steroids are initiated prior (10-20 minutes) or at least concomitant with the first dose of antibiotics *(2, 17)*. Therefore, innovative drugs that have combined antibacterial and anti-inflammatory functions to enable effective control over drug-resistant bacteria and inflammation-associated injuries at the earliest possible are critical for minimizing debilitation and mortality of infectious inflammation in the post-antibiotic era.

Human keratin 6a (K6a) is an intermediate filament protein highly expressed in various epithelial cells. We previously reported a direct antibacterial response of epithelial cells involving reorganization of the endogenous K6a filament network to upregulate the level of cytosolic K6a, which is processed by the ubiquitin-proteasome system to generate a series of short peptides. Importantly, the glycine-rich short peptides from the carboxyl terminal region of K6a (residues 515 to 559) termed keratin-derived antimicrobial peptides (KAMPs) are structurally flexible, salt-tolerant, and bactericidal via cell envelope disruptions and potentially intracellular targeting mechanisms *(18, 19)*. As such, KAMPs are naturally occurring, constitutive as well as inducible, and the only human example among the structurally unique non-αβ class of host defense peptides *(19, 20)*. Given that antimicrobial peptides often employ multiple bactericidal mechanisms including rapid cell lysis, the likelihood of bacterial resistance development or transmission is admittedly lower compared to the case of conventional antibiotics *(21-23)*. Moreover, a number of studies have showed that natural antimicrobial peptides can be modified by amino acid substitution to reduce cytotoxicity, enhance antimicrobial activity and spectrum, or incorporate anti-biofilm or immunomodulatory functions *(24, 25)*, demonstrating the feasibility of developing, optimizing, and tailoring antimicrobial peptide-based treatments for specific medical conditions.

In view of the outstanding potential of host defense peptides as new anti-infective alternatives, experimental animal models of skin and lung infections have been commonly used to examine their bacterial clearance efficacy per se. On the other hand, the therapeutic potential of engineered peptides in immune regulation was often assessed in animal models of sterile inflammation, in particular endotoxin-induced sepsis, which focus on mortality rate when evaluating treatment outcomes. As such, there is a paucity of preclinical studies aiming to reduce tissue damage and debilitating functional loss especially by concomitant control of infection and inflammation with bifunctional agents. Here, we used primary mouse neutrophils and macrophages, and murine models of sterile corneal inflammation, to first reveal KAMPs’ anti-inflammatory function, which is independent from their antimicrobial activity. We then examined their mechanisms of inflammation suppression and showed that, unlike many antimicrobial peptides that interact and neutralize bacterial ligands (LPS and LTA), KAMPs directly bind to TLR2, CD14 and MD-2 to block docking of these inflammatory molecules. KAMPs binding also promotes bacterial ligand-free TLR2 and TLR4 endocytosis, thus reduces the cell surface availability of these receptors for inflammatory activation. Notably, KAMPs do not trigger TLR signaling that activates NF-кB and IRF3 as LPS and LTA do. Furthermore, we demonstrated the therapeutic potential of bifunctional KAMPs in prevention and treatment of Gram-negative *(P. aeruginosa)* and Gram-positive *(S. aureus)* bacterial keratitis in a murine model, as evidenced by their effectiveness in concomitant control of infectious load, neutrophil and macrophage recruitment, and inflammation-associated corneal opacification. Also, unlike corticosteroids, KAMPs accelerate - as opposed to depress - corneal re-epithelialization while inflammation is alleviated. This study highlights the multiple intervening mechanisms of KAMPs in TLR activation, and demonstrates their potential therapeutic benefits for prevention of tissue damage and severe debilitation via concomitant control of infection and aggressive inflammation.

## RESULTS

### KAMPs are non-cytotoxic, non-apoptotic, non-hemolytic and non-barrier disruptive

To begin evaluating therapeutic potential of KAMPs, we tested whether these peptides pose undesirable risks for negative effects on cell viability. It has been shown that KAMP-19 (RAIG<u>GGLSSVGGGS</u>STIKY) at 200 μg/ml is non-toxic to cultured human corneal epithelial (hTCEpi) cells while protecting the cells against bacterial invasion and bacteria-induced cytotoxicity in a 3-h in vitro assay *(18)*. Here, we lengthened the incubation time to 24 h for hTCEpi cells *(26)* to be exposed to two truncated derivatives of KAMP-19, i.e. KAMP-18C (RAIG<u>GGLSSVGGGS</u>STIK) and their core sequence KAMP-10 (<u>GGLSSVGGGS</u>). These two peptides were chosen based on our previous report that characterized their molecular structures and bactericidal mechanisms *(19)*. We found that the levels of lactate dehydrogenase (LDH) released into the media from dead cells were comparable to the baseline (untreated cells) at all concentrations tested (up to 200 μg/ml), indicating that KAMP-10 and KAMP-18C did not induce cell death (fig. 1A). Further testing was performed to determine whether KAMPs induce more subtle changes to the viability of corneal epithelial cells. By detecting apoptotic cells that are undergoing DNA fragmentation in a standard TUNEL assay, we found that both KAMPs did not induce apoptosis during the 24-h incubation period (fig. 1B). While KAMPs appeared to be safe to corneal epithelial cells, we examined their effects on the general barrier integrity of corneal epithelium. Intact mouse eyeballs incubated with KAMP-10 or KAMP-18C ex vivo for 3 h did not show fluorescein penetration into the corneas (fig. 1C), drastically opposing to those subject to tissue paper blotting, a procedure that has been shown to compromise corneal epithelial barrier function *(27)*. Furthermore, we assessed potential hemolytic effects of KAMPs and found that the level of hemoglobin released from any lysed red blood cell treated with KAMP-10 or KAMP-18C was comparable to the baseline of untreated samples (fig. 1D). Taken together, the results support a favorable safety profile for KAMPs.

**Fig. 1.**
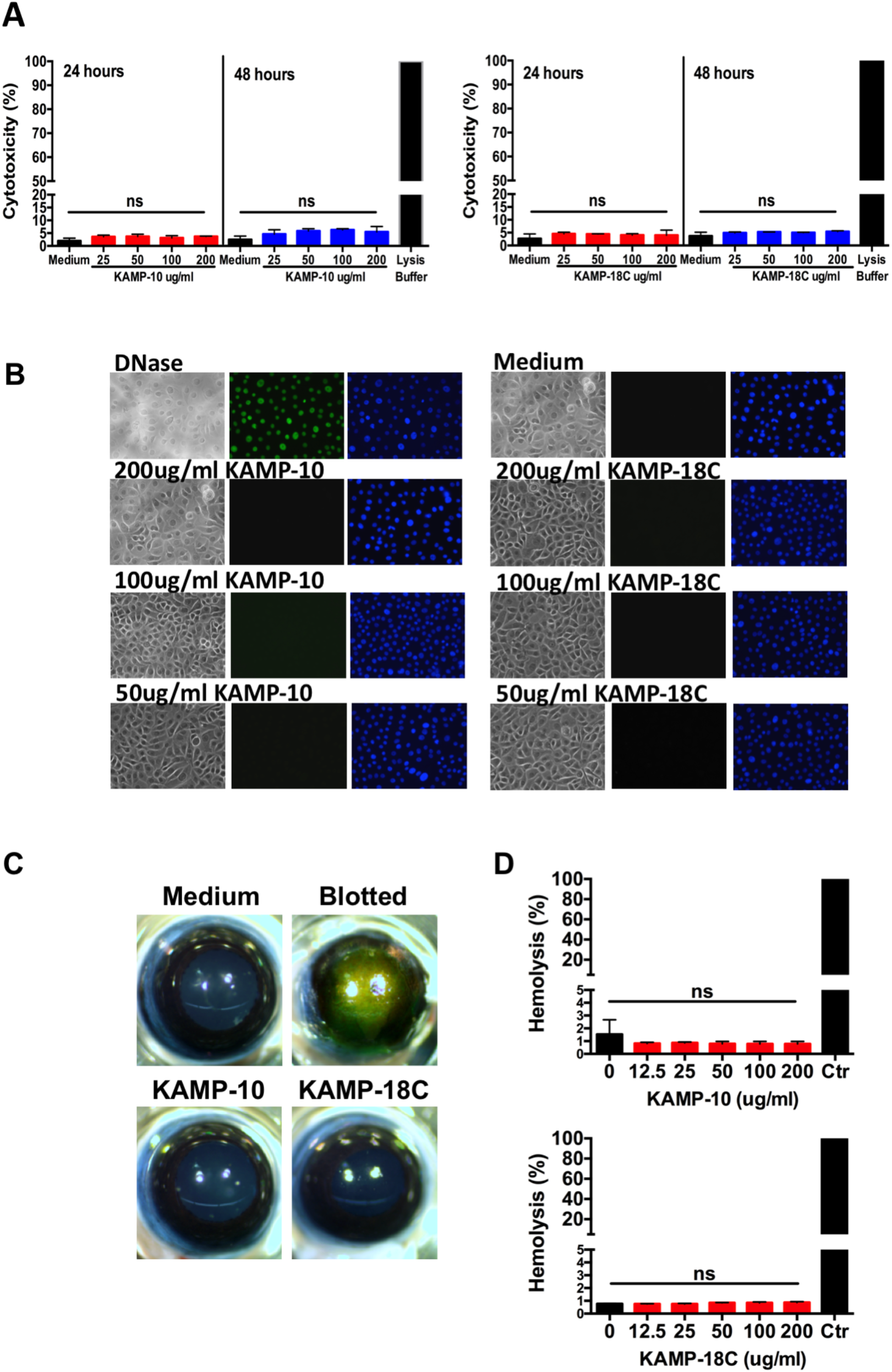
KAMP-10 and KAMP-18C are non-cytotoxic. (**A**) Human corneal epithelial cells (hTCEpi) were incubated with KAMP-10 or KAMP-18C at the indicated concentrations for 24 or 48 hours. Mock-treated cells and those treated with Triton-X lysis buffer were used as baseline and positive control (100% cytotoxicity), respectively. LDH activities in cell culture media were assayed using the colorimetric LDH Cytotoxicity Kit and normalized to the positive control to determine % cytotoxicity. *P* > 0.05 (+peptide vs. medium, Student’s t-test). (**B**) Apoptosis of hTCEpi cells was assayed by the fluorescent TUNEL Assay Kit after 24-hour incubation with the indicated concentration of KAMPs. DNase treatment was used as positive control for apoptosis (green nuclei). Cells were counterstained with DAPI (blue nuclei) before imaging with a phase contrast (cell morphology) and fluorescence microscope (20x objective). (**C**) Enucleated mouse eyes were rinsed with PBS, then submerged in 1 mg/ml of KAMP-10 or KAMP-18C for 3 hours at 35°C, followed by fluorescein staining to assess corneal barrier integrity. Intact corneas are impermeable to the stain. Corneas that were tissue paper blotted to disrupt epithelial barrier function were used as the positive control of fluorescein staining. (**D**) Human red blood cells were incubated with the indicated concentration of KAMP-10 or KAMP-18C or mock treated for 3 hours. The levels of free heme released to the media from lysed cells were measured then normalized to cells that were treated with Triton-X (complete lysis). Mean ± SD (n=3) are shown. Non-significant (ns): *P* > 0.05 (ANOVA).

### KAMPs suppress the production of bacterial ligand-induced proinflammatory and chemotactic cytokines from primary murine neutrophils and macrophages

To assess the immunomodulatory activity of KAMPs, freshly isolated mouse bone marrow neutrophils and resident peritoneal macrophages were stimulated with two common inflammatory bacterial components - purified Gram-negative *P. aeruginosa* LPS and Gram-positive *S. aureus* LTA (500 ng/ml) - in the presence or absence of KAMPs (50 to 200 μg/ml) for 18 h, followed by quantitative measurements of secreted cytokines in the culture supernatants by ELISA. We found that KAMP-10 and KAMP-18C demonstrated dose-dependent inhibitory effects on induced secretion of proinflammatory CXCL1, CXCL2, CXCL10, IL-6, TNF*α* and G-CSF from both neutrophils (fig. 2 and S1; A and B) and macrophages (fig. 2 and S1; C and D). Notably, KAMP-10 exerted a more potent anti-inflammatory effect against LPS-induced cytokine secretion (fig. 2 and S1; A and C), whereas KAMP-18C showed a more pronounced suppressive effect to LTA stimulation (fig. 2 and S1; B and D). As such, when the peptide treatment was applied 30 minutes before stimulation onset, we observed that the concentration of KAMP-10 that achieved complete inhibition against LPS induction, or KAMP-18C against LTA induction, was 100 μg/ml. When the peptide treatment was applied 30 minutes after stimulation onset, a higher dose (200 μg/ml) was needed to achieve comparable suppressive effects as seen in the pretreatment regime.

**Fig. 2.**
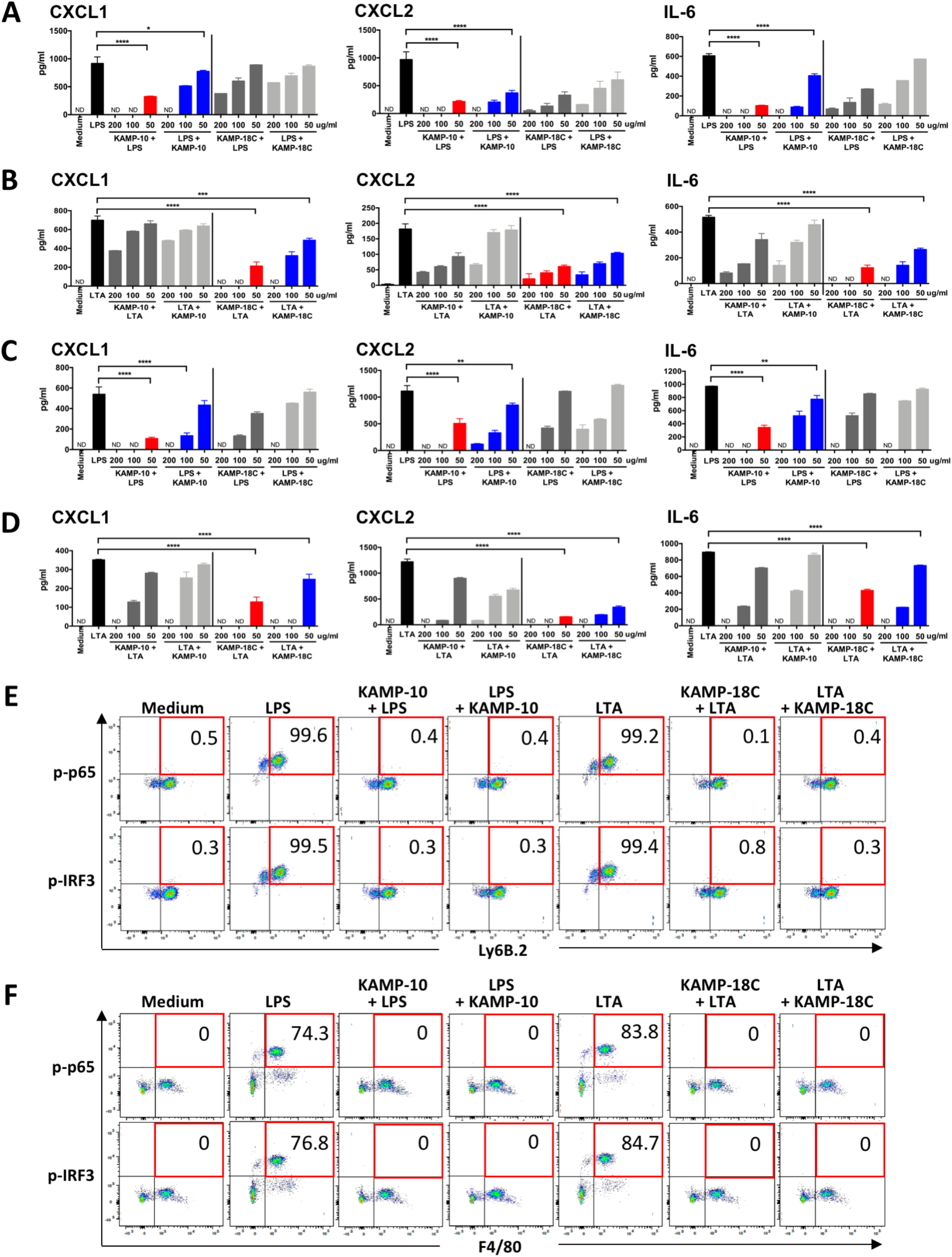
KAMP-10 and KAMP-18C attenuate LPS-and LTA-induced inflammatory responses in murine neutrophils and macrophages. (**A-D**) Enriched murine bone marrow neutrophils (A, B) and resident peritoneal macrophages (C, D) were stimulated with LPS (A, C) or LTA (B, D) (500 ng/ml) in the absence or presence of KAMP-10 or KAMP-18C at the indicated concentrations for 24 hours. The peptides were applied either 30 minutes before (red and dark grey) or 30 minutes after (blue and light grey) the stimulation. Cells mock-treated with medium served as baseline controls. CXCL1, CXCL2, IL-6 levels in culture supernatants were measured by ELISA. Mean ± SD (n=3 replicates) are shown. ND=non-detected. \**P* < 0.05, \*\**P* < 0.01, \*\*\**P* < 0.001, \*\*\*\**P* < 0.0001 (ANOVA with Dunnett’s post hoc test). The lowest concentration of KAMP yielding statistical significance are shown. See fig. S1 for additional three cytokines (CXCL10, TNF*α*, G-CSF). (**E-F**) Representative plots showing thioglycollate-elicited and IFN*γ*-treated peritoneal neutrophils (E) and macrophages (F) that were untreated or treated with LPS (in the presence or absence of 200 μg/ml KAMP-10) or LTA (in the presence or absence of 200 μg/ml KAMP-18C). Intracellular staining of phosphor-NFкB p65 (top) and phosphor-IRF3 (bottom) were analyzed by flow cytometry. Numbers at top right corners represent percentage of Ly6B.2^+^ neutrophils or F4/80^+^ macrophages that were p-p65^+^ or p-IRF3^+^.

Using multiplex immunobead assays, we further examined the suppressive effects of KAMP-10 and KAMP-18C on a wide range of proinflammatory and chemotactic cytokines produced by mouse bone marrow neutrophils and resident peritoneal macrophages. As complete abolition of cytokine production may not be desirable or necessary in most clinical situations, we chose a sub-effective dose of KAMP-10 and KAMP-18C to test against LPS and LTA respectively to evaluate their capacity of suppression. In the case of neutrophils (fig. S2), pretreatment with a low dose (50 μg/ml) of KAMPs significantly suppressed eleven LPS- or LTA-induced cytokines and chemokines. In comparison, post-treatment with the same dose of KAMPs generally yielded weaker suppression of these proinflammatory and chemotactic cytokines; yet IFNγ, IL-1β, CXCL1, CXCL2, CCL2 and G-CSF were still suppressed by more than 50%. In the case of macrophages (fig. S3), pretreatment with the low dose (50 μg/ml) of KAMPs significantly suppressed nine LPS- or LTA-induced cytokines and chemokines. Similar to neutrophils, post-treating macrophages with the same dose of KAMPs suppressed IFNγ and CCL2 secretion by more than 50%, in addition to CXCL10, CCL3, CCL4 and CCL5. Taken together, KAMPs exert potent anti-inflammatory effects on LPS- and LTA-stimulated neutrophils and macrophages.

Next, we utilized flow cytometry to confirm the attenuation effects of KAMPs on LPS- or LTA-induced NFкB and IRF3 signaling. As expected, the percentage of phospho-NFкB p65 or phospho-IRF3 expressing neutrophils (fig. 2E) and macrophages (fig. 2F) were substantially increased upon stimulation with LPS or LTA. In agreement with the suppression of inflammatory cytokine responses, treatment with KAMP-10 or KAMP-18C (200 μg/ml), whether it was initiated before or after the bacterial ligand stimulation, reduced the percentage of activated cells to the level of baseline (unstimulated cells).

### KAMPs suppress LPS- and LTA-induced acute corneal inflammation in mice

Next, to examine in vivo relevance of the cytokine-suppressive activity of KAMPs, we tested the effects of KAMP-10 and KAMP-18C on the early inflammatory response in abraded mouse corneas challenged with LPS (fig. 3, A and B) or LTA (fig. 3, C and D) for 24 h. As expected, LPS and LTA alone elevated CXCL1, CXCL2, CXCL10, TNF*α*, IL-6 and G-CSF levels in the corneas compared with trauma controls that were treated with PBS. In contrast, one dose of KAMP-10 topically applied 30 min before or after LPS challenge reduced levels of these cytokines to those of the trauma control (fig. 3A). Similar degree of suppression by KAMP-18C on corneal cytokine levels was also observed in LTA-challenged mice (fig. 3C). To determine whether KAMPs suppress LPS- or LTA-induced acute inflammatory cell recruitment to corneas, we quantified total leukocytes (CD45^+^), neutrophils (CD45^+^Ly6B.2^+^F4/80^-^) and macrophages (CD45^+^F4/80^+^) in dissected whole corneas at 24 h post-challenge by flow cytometry. In sync with reduced proinflammatory and chemotactic cytokines in corneas, KAMP-10 treatment applied before or after LPS challenge significantly reduced numbers of total infiltrating leukocytes (by 86 and 97% respectively), including neutrophils (by 89 and 97%) and macrophages (by 77 and 95%) (fig. 3B). These results were supported by the significant reduction of immunofluorescence staining of Ly6B.2-expressing cells by KAMP-10 treatment in LPS-challenged mouse corneas (fig. S4). Consistently, KAMP-18C treatment before or after LTA challenge also reduced recruitment of total leukocytes (by 86 and 94% respectively), neutrophils (by 86 and 92%) and macrophages (by 88 and 96%) (fig. 3D). Taken the ex vivo experiments using purified myeloid cells (fig. 2) together with the in vivo studies using a mouse model of sterile corneal inflammation (fig. 3), the data indicated that KAMPs have potent anti-inflammatory activity that is independent of their previously characterized antimicrobial function *(18, 19)*.

**Fig. 3.**
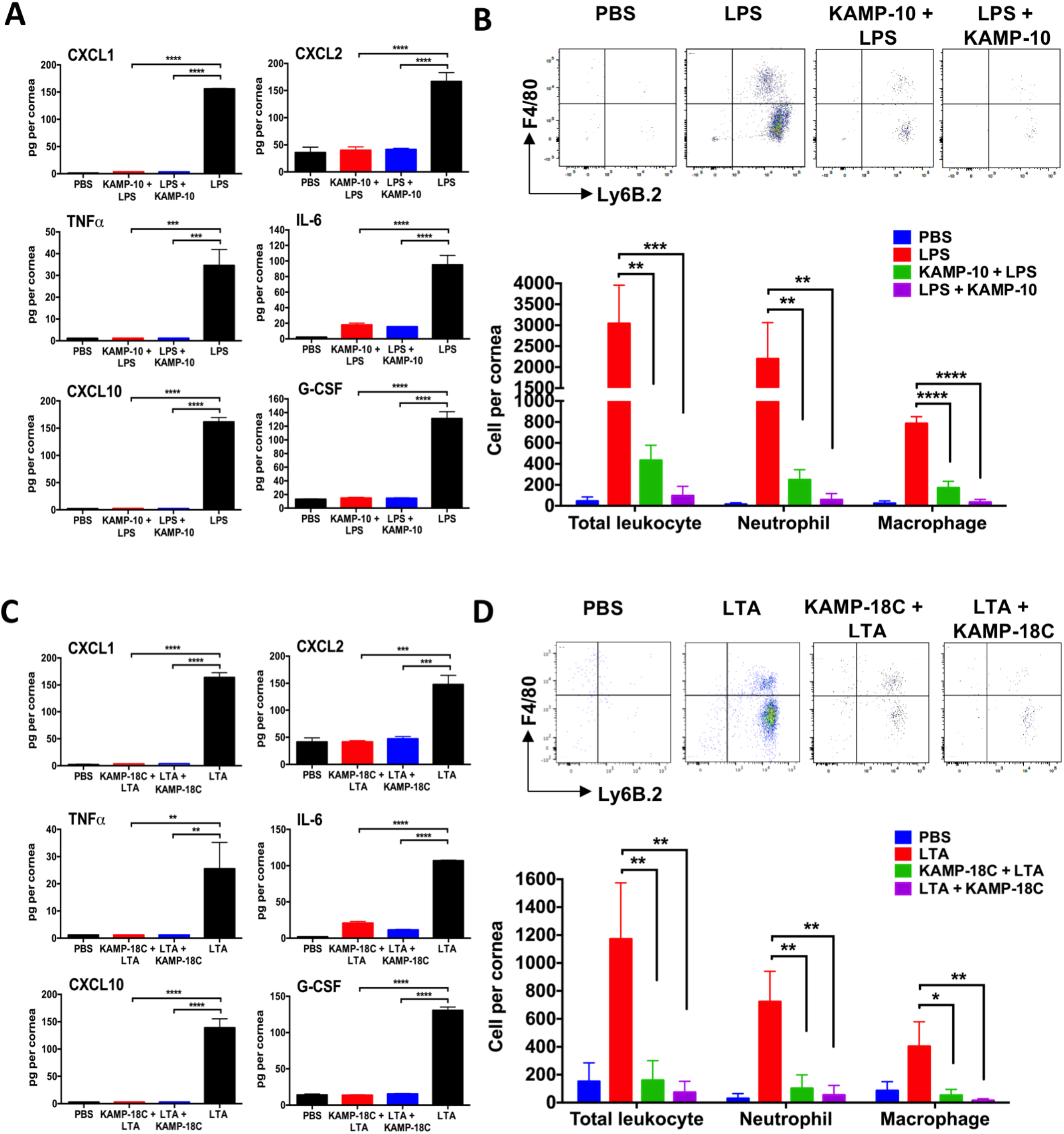
KAMP-10 and KAMP-18C suppress LPS- and LTA-induced corneal inflammation, respectively, in a murine model. (**A-B**) Purified *P. aeruginosa* LPS or (**C-D**) *S. aureus* LTA (20 μg) or PBS only was topically applied to scarified mouse corneas. The eyes were untreated, or treated with one topical application of KAMP-10 or KAMP-18C either 30 minutes before (100 μg/ml KAMP-10/18C + LPS/LTA) or 30 minutes after stimulation (LPS/LTA + 200 μg/ml KAMP-10/18C). After 24 hours, mice were euthanized and whole corneas were dissected for analysis. (A, C) Amounts of cytokines per cornea were determined by ELISA. (B, D) Representative FACS plots showing Ly6B.2 and F4/80 gated cells among total CD45^+^ cells (top). Total leukocytes (CD45^+^), neutrophils (CD45^+^Ly6B.2^+^F4/80^-^) and macrophages (CD45^+^F4/80^+^) per cornea were quantified (bottom). Mean ± SD (n=3 mice) are shown. \**P* < 0.05, \*\**P* < 0.01, \*\*\**P* < 0.001, \*\*\*\**P* < 0.0001 (ANOVA with Dunnett’s post hoc test).

### KAMPs block LPS and LTA activation via direct binding to MD-2 and TLR2/CD14

We next questioned how KAMPs dampen the cytokine responses of LPS- or LTA-stimulated neutrophils and macrophages. Since our previous report demonstrated low to moderate binding affinities of KAMP-10 to LPS and of KAMP-18C to LTA *(19)*, we anticipated that KAMPs exert anti-inflammatory effects via multiple routes apart from the typical neutralization mechanism shared among various antimicrobial peptides *(28, 29)*. To test the hypothesis that direct binding of KAMP-10 to TLR4/MD-2, and KAMP-18C to TLR2/CD14, inhibits their respective activation by LPS and LTA, we first conducted molecular docking simulations to explore potential interactions between these receptors and KAMPs. As shown in fig. 4A, KAMP-10 could be confined in the hydrophobic barrel-like cavity of MD-2 in both soluble form (sMD-2) and TLR4-associated form (MD-2/TLR4) similarly to LPS (the acyl chains of lipid A core) *(30-33)*. KAMP-18C was predicted to interact with the hydrophobic pockets at the TLR2 central domain (fig. 4B) and the CD14 N-terminal domain (fig. 4C), which both have been characterized as the binding sites for acylated ligands including LTA *(34-37)*.

**Fig. 4.**
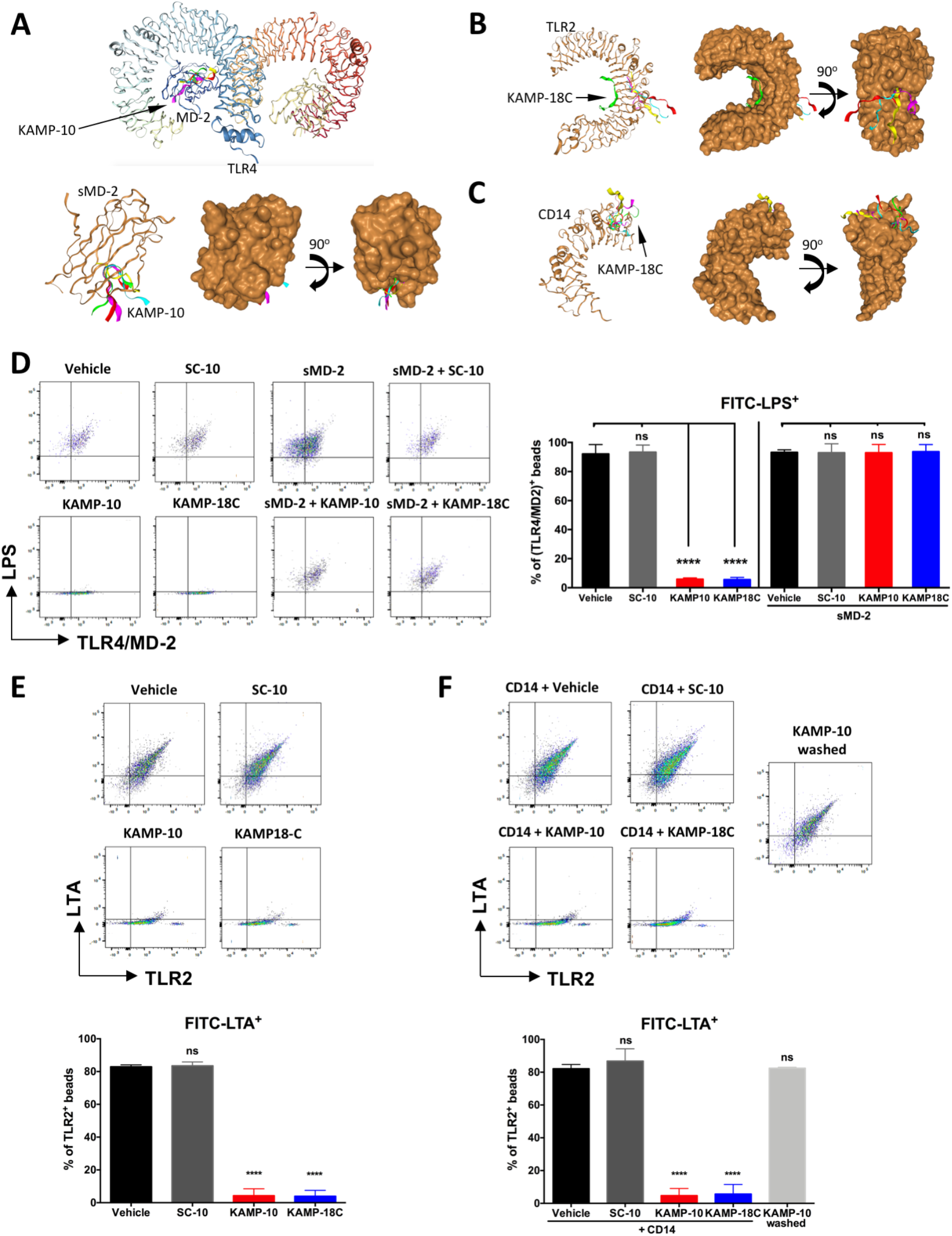
KAMP-10 couples with MD-2, and KAMP-18C with TLR-2 and CD14, to disrupt LPS/MD-2/TLR4 and CD14/LTA/TLR2 interactions. (**A-C**) Molecular docking of KAMP-10 and KAMP-18C (ribbon representation) to TLRs and their co-receptors (ribbon and surface representations). Top five models of the peptides (1 to 5: yellow, green, red, magenta, cyan) docked to the receptors are shown. (A) KAMP-10 was predicted to bind to the hydrophobic cavity of MD-2 that is in complex with TLR4 (top) or in the soluble form (bottom). (B-C) KAMP-18C was predicted to bind to hydrophobic pocket at the central domain of TLR2 (B) and at the N-terminal domain of CD14 (C). (**D**) Representative FACS plots (left) and quantification (right) showing the extent of LPS binding to TLR4/MD-2-conjugated beads. Latex beads with human TLR4-MD-2 complex conjugated on the surface were incubated with the peptides (KAMP-10 or SC-10 at 48 μg/ml, or KAMP-18C at 96 μg/ml) in the absence or presence of soluble MD-2 (1 μg/ml), washed, then incubated with FITC-LPS (1 μg/ml). Bead surface TLR4 was labeled with antibody followed by flow cytometry. Percent of TLR4-MD-2^+^ beads that were FITC-LPS^+^ is shown. (**E-F**) Representative FACS plots (top) and quantification (bottom) showing the extent of LTA binding to TLR2-conjugated beads. (E) Human TLR2-conjugated beads were incubated with the peptides (48 or 96 μg/ml), washed, then incubated with LTA (1 μg/ml) and CD14 (0.1 μM). (F) CD14 (0.1 μM) was preincubated with equal molar of peptides (0.1 μM) before TLR2-conjugated beads and LTA (1 μg/ml) were added. In comparison, KAMP-10 alone (0.1 μM) was preincubated with TLR2-conjugated beads, followed by bead washing then incubation with LTA (1 μg/ml) and CD14 (0.1 μM). TLR2 and LTA were labeled with antibody for flow cytometric analysis. Percent of TLR2^+^ beads that were FITC-LTA^+^ is shown. Mean (n=3 replicates) ± SD. \**P* < 0.05, \*\**P* < 0.01, \*\*\**P* < 0.001, \*\*\*\**P* < 0.0001 (ANOVA with Dunnett’s post hoc test).

Given that KAMPs could potentially block the ligand binding sites of TLRs and their co-receptors, we used latex beads that were surface-conjugated with either TLR4/MD-2 complex or TLR2 to examine the effects of KAMPs on LPS and LTA binding. As shown by flow cytometric analyses, preincubation of TLR4/MD-2 complex-conjugated beads with KAMP-10 or KAMP-18C, but not the scrambled KAMP-10 variant (SC-10), inhibited subsequent LPS binding (fig. 4D). This inhibitory effect was abolished when soluble MD-2 was present during preincubation (fig. 4D), indicating that KAMP-10 and KAMP-18C compete with LPS for the same binding site on MD-2 and thus prevents TLR4/MD-2/LPS complex formation. Likewise, we found that preincubation of TLR2-conjugated beads with KAMP-10 or KAMP-18C prevented subsequent CD14-mediated binding of LTA to TLR2 (fig. 4E), indicating that these peptides directly block the LTA-binding site on TLR2. In another setting when CD14 was preincubated with equal molar of KAMP-10 and KAMP-18C (peptide concentration now equivalent to 600-fold dilution of the one used in aforementioned experiments) before exposure to TLR2-conjugated beads and LTA, LTA binding to TLR2 was abolished (fig. 4F). Notably, in the absence of CD14, preincubation with KAMP-10 at this low concentration was not sufficient to block subsequent CD14-mediated LTA binding to TLR2 (fig. 4F). Together, the data strongly suggested that KAMPs can disrupt the interaction between CD14 and LTA and thus prevent CD14 from transferring the bacterial ligand to TLR2. It is most likely that CD14 promotes KAMP-10 and KAMP-18C transfer to TLR2 to enhance their direct blocking effects against LTA binding.

### KAMPs induce endocytosis of bacterial ligand-free TLR4 and TLR2

In addition to cell surface activation of TLR proinflammatory signaling, bacterial components complexed with TLR2 (e.g. LTA) or TLR4/MD-2 (e.g. LPS), together with CD14, are internalized to induce endosomal signaling that activates interferon (IFN) regulatory factors (IRFs) for type I IFN gene expression *(39, 40)*. TLR endocytosis is considered as an important negative regulatory mechanism for controlling cell surface availability of TLRs and thus the magnitude of pro-inflammatory responses *(41-44)*. To examine whether KAMPs modulates TLR endocytosis, thioglycollate-elicited mouse peritoneal neutrophils and macrophages were treated with LTA, LPS or KAMPs alone, followed by flow cytometric analyses of extracellular and intracellular TLRs. Similar to LPS, incubation with KAMP-10 and KAMP-18C but not SC-10 alone for 1 h reduced cell surface staining but increased intracellular staining of TLR4/MD-2 for both neutrophils (fig. 5A) and macrophages (fig. 5C). We also observed that KAMP-10 and KAMP-18C but not SC-10 alone triggered internalization of TLR2 as LTA does (fig. 5, B and D). These findings, together with those demonstrating the ability of KAMPs to engage MD-2, CD14 and TLR2 (fig. 4), supported the notion that the anti-inflammatory mechanisms of KAMPs are multifaceted, including direct blocking of CD14, TLR2 and MD-2 to prevent recognition of LTA and LPS by cell surface TLR2 and TLR4, as well as direct induction of receptor internalization before they are engaged by these bacterial ligands.

**Fig. 5.**
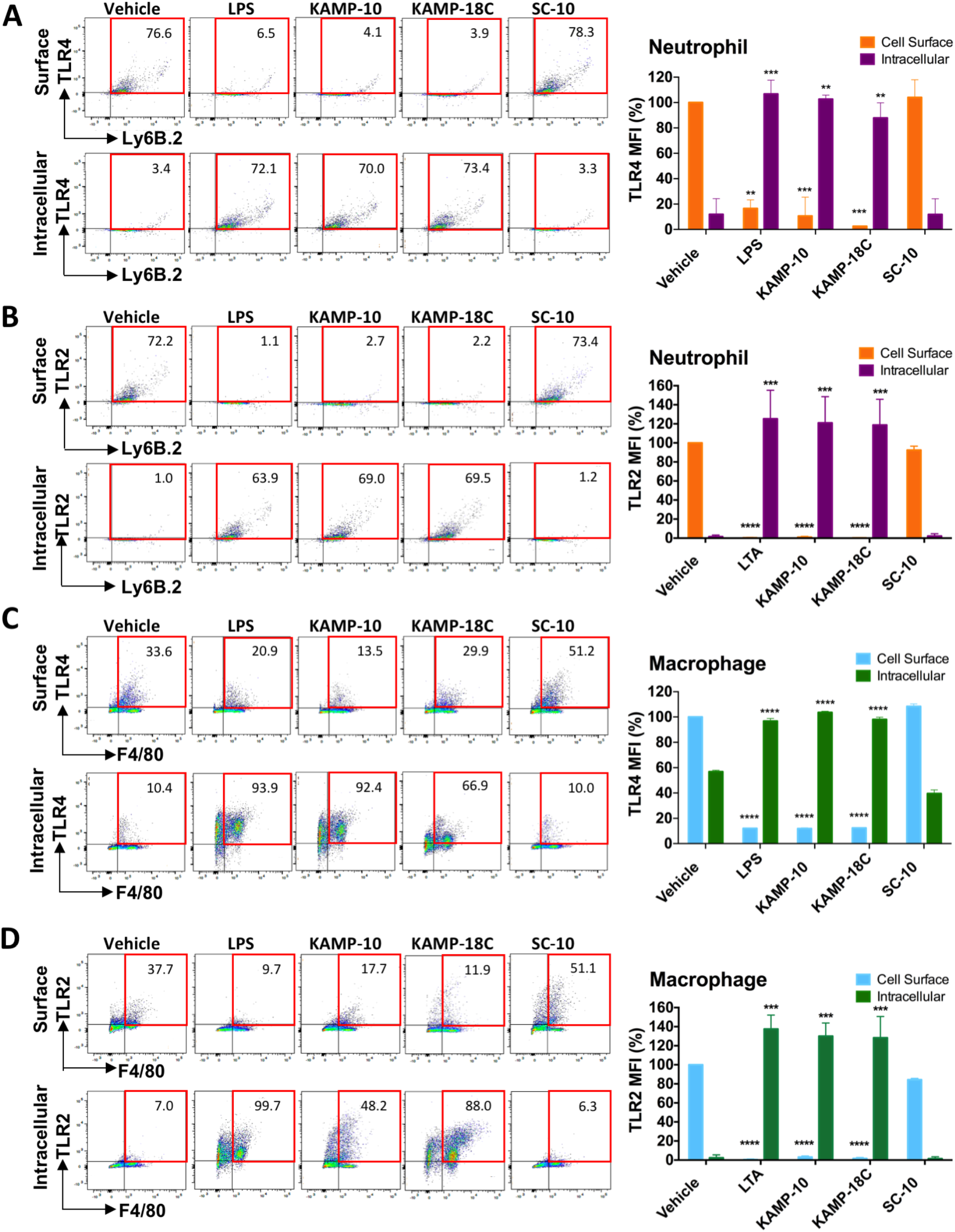
KAMP-10 and KAMP-18C induce endocytosis of bacterial ligand-free TLR4 and TLR2 respectively. Thioglycollate-elicited (**A-B**) mouse peritoneal neutrophils and (**C-D**) macrophages were unstimulated or stimulated with LPS (A, C) or LTA (B, D) (500 ng/ml), or KAMP-10, KAMP-18C or SC-10 (200 μg/ml) alone for 1 hour. Cell surface and intracellular levels of TLR4 and TLR2 were detected by flow cytometry. Numbers at top right corners of representative flow cytometry plots (left) are percentage of Ly6B.2^+^ neutrophils or F4/80^+^ macrophages that were positively stained for TLR4 or TLR2. Median fluorescence intensity (MFI) of cell surface and intracellular TLR4 and TLR2 were also quantified (right). Mean (n=3 replicates) ± SD are shown. \*\**P* < 0.01, \*\*\**P* < 0.001, \*\*\*\**P* < 0.0001 (same compartment comparison between treatment groups and vehicle; ANOVA with Dunnett’s post hoc test).

### Topical KAMPs prevent bacterial infection and inflammation in a murine keratitis model

Considering the therapeutic potential of coupled antimicrobial and anti-inflammatory activities for simultaneous abrogation of infection and the accompanying tissue damage, we first evaluated the efficacy of bifunctional KAMPs for the prevention of infection and inflammation in a murine bacterial keratitis model. As shown in fig. 6, mouse corneas were scarified and immediately inoculated with a live Gram-negative bacterium (an amikacin-resistant spontaneous mutant of *P. aeruginosa* keratitis isolate) or Gram-positive bacterium (a standard laboratory strain of *S. aureus)* with or without concomitant topical administration of KAMPs, and disease progression was monitored for 48 h. Beginning at 24 h after infection, corneal opacification and irregularity were prominent in infected mice without prophylactic treatment, whereas the two groups given KAMP-10 (against *P. aeruginosa)* (fig. 6, A and B) or KAMP-18C (against *S. aureus)* (fig. 6, E and F) appeared normal with slight surface irregularity only. And as expected, corneal pathology continued to progress in unprotected mice, contrasting the healthy-looking eyes of the two groups of KAMP-treated mice (mean clinical score 0) at 48 h after infection. Consistent with the disease presentations, high bacterial loads (mean 10^4^-10^5^ CFU) were found in all unprotected mouse eyes, whereas the KAMPs-treated eyes were free of bacteria (fig. 6, C and G). Not only did the prophylactic administration of KAMPs during bacterial inoculation eliminate pathogens (including the antibiotic-adapted mutant), it also prevented massive cellular infiltration, a normal host response triggered by microbial invasion. As shown by flow cytometric analysis (fig. 6, D and H), both groups of KAMP-treated mice had significantly lower numbers of total leukocytes (CD45^+^), neutrophils (CD45^+^Ly6B.2^+^F4/80^-^) and macrophages (CD45^+^F4/80^+^), in the whole corneas at 48 h post-infection compared with the unprotected group. Since antibiotic-killed *P. aeruginosa* and *S. aureus* (non-infectious and yet possessing inflammatory ligands) can still trigger a high magnitude of neutrophil recruitment in the same model of murine keratitis *(46, 47)*, the observed immunosuppressive effects of prophylactic KAMP-10 and KAMP-18C could not be attributed solely to effective bacteria killing. Instead, they are consistent with the primary effects of these peptides on sterile inflammation induced by LPS and LTA (fig. 2 and 3). These findings suggested that topical administration of KAMP-10 or KAMP-18C on wounded (susceptible) corneas provides more effective prophylactic protection than antibiotics alone against both bacterial infection and associated inflammation in subjects at risk.

### Topical KAMPs concurrently abrogate bacterial infection and inflammation-associated tissue damage in a murine keratitis model

We next assessed the potential of KAMPs as a treatment option for managing bacterial infections of the cornea, in which the acute proinflammatory responses must also be reduced to avoid tissue damage and vision loss. Instead of concomitant administration of KAMPs with bacterial inoculum previously conducted in the prophylaxis study, here we treated ongoing *P. aeruginosa* keratitis with KAMPs, starting one day after bacterial inoculation when the disease had already been established. As poor absorption is a major concern for ocular drugs *(48)*, KAMPs were mixed with a recognized carrier - liposome *(49, 50)* - immediately before topical administration to enhance peptide delivery into corneal stroma, where bacteria and immune cell infiltrates are accumulating. As shown in fig. 7 (A-E), mouse corneas infected by amikacin-resistant *P. aeruginosa* showing similar disease severity one day after bacterial inoculation (mean clinical score 5.3-5.5, p > 0.05) were topically treated with liposomes alone (carrier control), liposomes/KAMP-10 mixture or liposomes/KAMP-18C mixture three times a day for 3 days. We observed that both groups of KAMP-treated eyes had substantially lower disease scores each day compared with the infected eyes that received carriers only (fig. 7, A and B). In relation to pre-treatment state, mice received KAMP-10 began to show marked clinical improvement after two days of treatment (fig. 7B). KAMP-18C treatment also improved disease outcome of *P. aeruginosa* keratitis over the 3-day course, although the reduction of disease scores was not statistically significant. Consistent with the disease presentations, flow cytometric analysis of acute inflammatory cells in the infected mouse corneas on day 3 post-treatment showed that the recruitment of total leukocytes, neutrophils and macrophages was substantially reduced by KAMPs (fig. 7, D and E). Importantly, the suppressed immune responses did not result in uncontrolled bacterial growth. Indeed, both KAMPs demonstrated significant contribution to bacterial eradication, as indicated by the 2-log reduction of *P. aeruginosa* in the eye on day 3 post-treatment compared with the carrier control group (fig. 7C). The results showed that topical KAMPs achieved simultaneous control of the Gram-negative bacterium and inflammation-associated tissue damage.

**Fig. 6.**
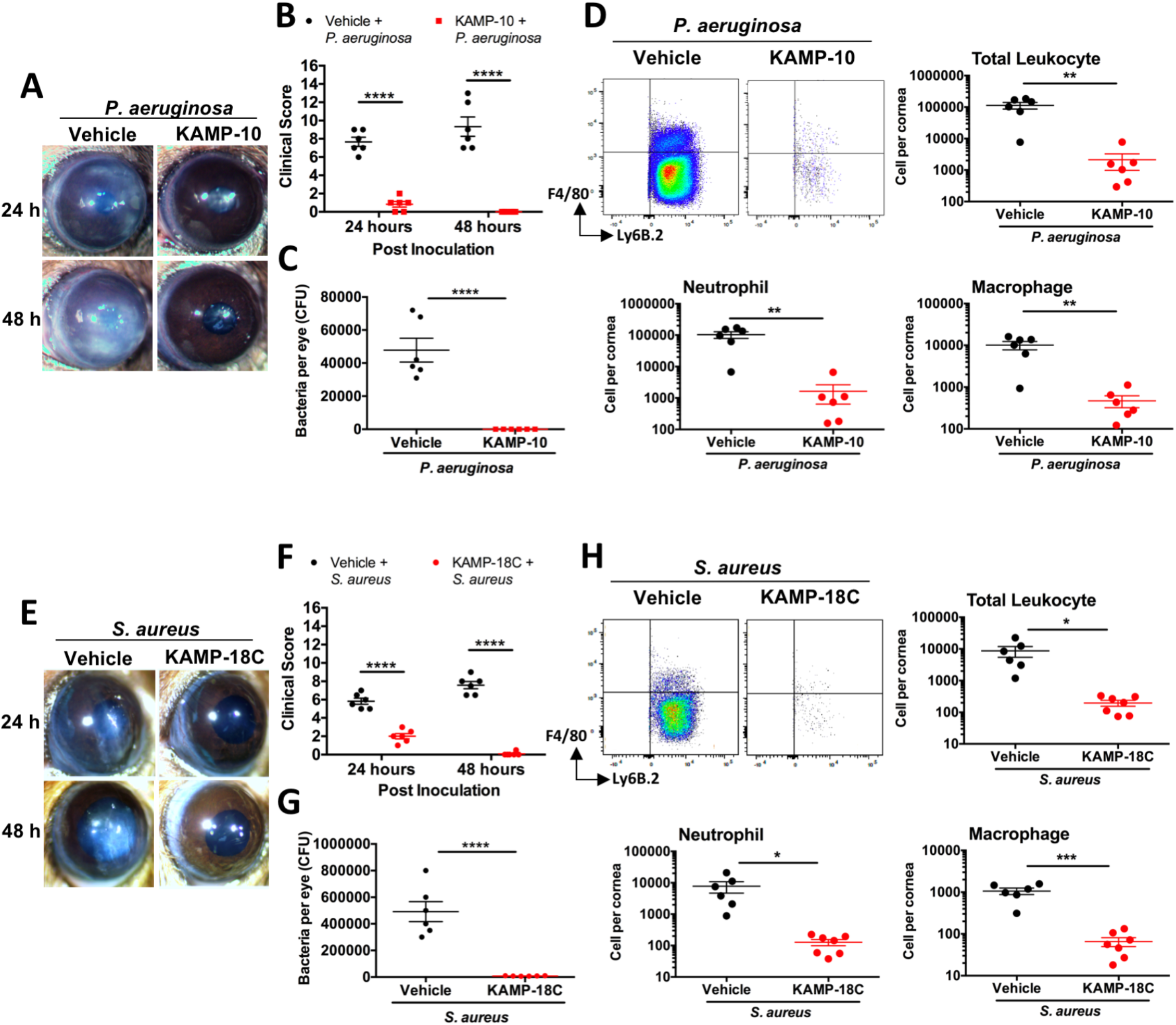
KAMP-10 and KAMP-18C prevent bacterial corneal infection in a murine model. Scarified mouse corneas were inoculated with (**A-D**) *P. aeruginosa* (10^5^ CFU) in the absence or presence of KAMP-10 (200 ng), or (**E-H**) *S. aureus* (10^6^ CFU) in the absence or presence of KAMP-18C (200 ng). Disease presentations at 24 and 48 h post-inoculation were photographed (A, E) and severity was scored corresponding to the extents of opacification and surface irregularity (B, F). At 48 h post-inoculation, mice were euthanized followed by quantification of bacterial load (C, G) and immune cells (D, H) in dissected corneas. (H) Representative FACS plots showing total CD45+ live cells from a cornea gated for Ly6B.2 and F4/80 markers. Leukocytes (CD45^+^), neutrophils (CD45^+^Ly6B.2^+^F4/80^-^) and macrophages (CD45^+^F4/80^+^) per cornea were quantified. Mean ± SEM (n=6-7 mice) are shown. \**P* < 0.05, \*\**P* < 0.01, \*\*\**P* < 0.001, \*\*\*\**P* < 0.0001 (Two-tailed Student’s t-test).

**Fig. 7.**
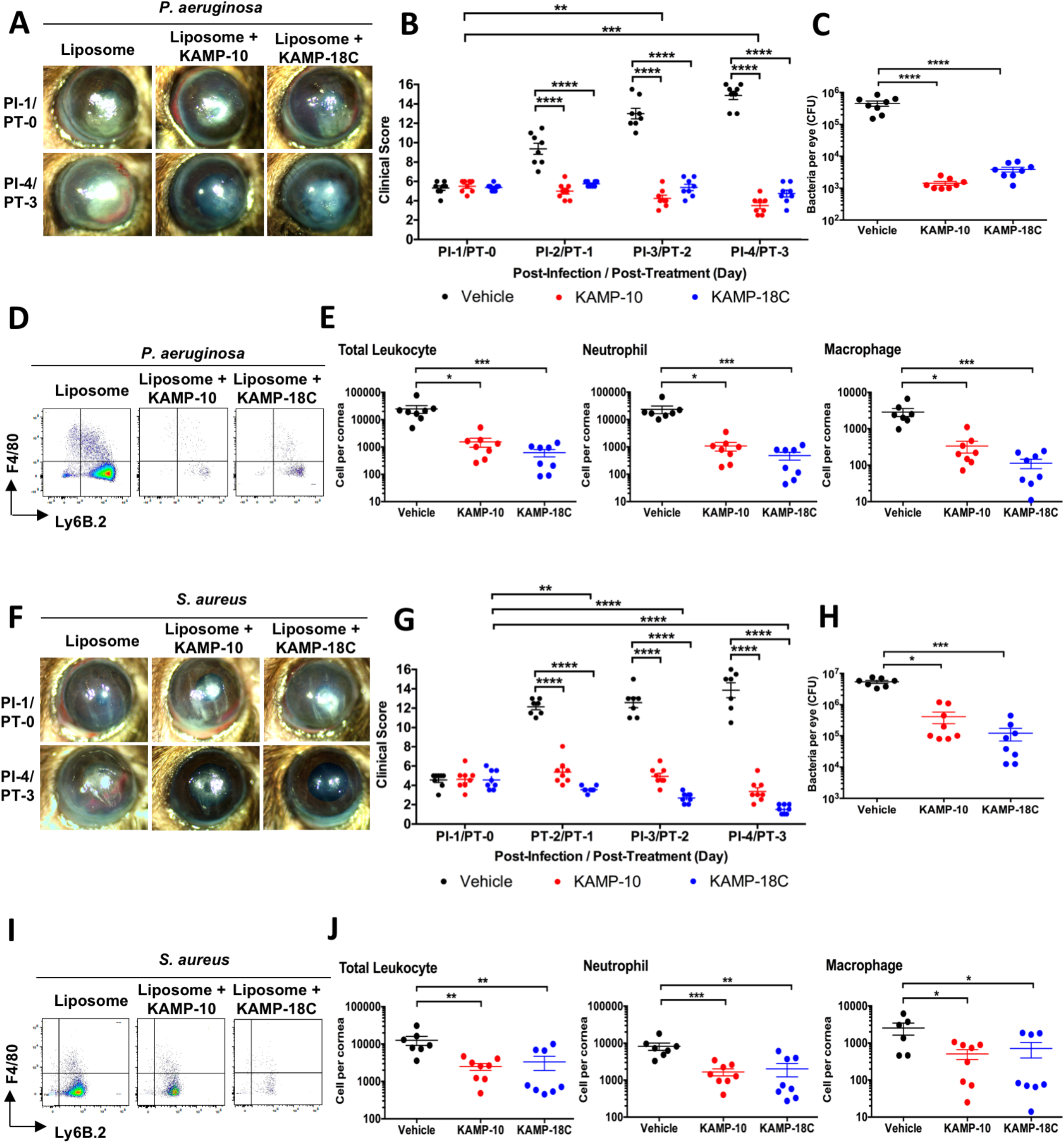
KAMP-10 and KAMP-18C effectively control bacterial burden and alleviate inflammation in a murine infectious keratitis model. Scarified mouse corneas were inoculated with (**A-E**) *P. aeruginosa* (10^5^ CFU) or (**F-J**) *S. aureus* (10^6^ CFU). At day 1 post-infection (PI-1), topical treatments (3 times daily for 3 days) with liposome alone or liposome-associated KAMP-10 or KAMP-18C (5 μg peptide) commenced. At day 1, 2, 3 and 4 post-infection, equivalent to day 0, 1, 2 and 3 post-treatment (PT), disease presentations were photographed (**A, F**) and scored corresponding to severity of opacification and surface irregularity (**B, G**). At day 4 post-infection, mice were euthanized followed by quantification of bacterial load (**C, H**) and immune cells (D-E, I-J) in dissected corneas. (**D, I**) Representative FACS plots showing total CD45+ live cells from an infected cornea gated for Ly6B.2 and F4/80 markers. (**E, J**) Leukocytes (CD45^+^), neutrophils (CD45^+^Ly6B.2^+^F4/80^-^) and macrophages (CD45^+^F4/80^+^) per cornea were quantified. Mean ± SEM (n=7-8 mice) are shown. \**P* < 0.05, \*\**P* < 0.01, \*\*\**P* < 0.001, \*\*\*\**P* < 0.0001 (ANOVA with Dunnett’s post hoc test (except E, H); Krustal-Wallis with Dunn’s post hoc test (E, H)).

To examine whether KAMPs are also effective against Gram-positive bacterial keratitis, we infected mouse corneas with *S. aureus* and commenced topical treatment with liposomes alone or liposome/KAMP mixtures one day after bacterial inoculation (fig. 7, F to J). Consistent with the findings from the *P. aeruginosa* keratitis model, *S. aureus* infected eyes treated with KAMPs had reduced pathological scores each day compared with the carrier-treated group (fig. 7, F and G). In relation to pre-treatment state, significant clinical improvement began one day following treatment with KAMP-18C (fig. 7G). Furthermore, both KAMPs effectively suppressed infiltration of total leukocytes, neutrophils and macrophages (fig. 7, I and J) while substantially reducing the infectious load (fig. 7H) in the *S. aureus*-infected corneas. This model of Gram-positive bacterial keratitis again demonstrated that concomitant control of infection and inflammation by the dual-function KAMPs is efficacious.

Since impaired would healing is a great concern over the use of immunosuppressive agents (e.g. corticosteroids) on compromised barriers, we examined the effects of KAMPs on corneal re-epithelialization in vivo. Mouse central corneas with the epithelium mechanically removed were topically treated with KAMP-10 or KAMP-18C alone. Consistent with the immunosuppressive effects of KAMPs on sterile LPS/LTA-induced inflammation (fig. 2 and 3), treatment with KAMPs yielded significant reduction of CXCL1, CXCL2, CXCL10, IL-6, TNF*α* and G-CSF proteins in the wounded corneas at 6 h post-injury as compared with the mock (vehicle) treatment group (fig. 8B). There were also fewer total leukocytes, particularly neutrophils and macrophages in the KAMPs-treated corneas at 24 h post-injury (fig. 8C). Despite that the early necessary inflammatory responses for wound healing was dampened, we found that wound closure was substantially accelerated by the treatment with KAMPs, as evidenced by greater reductions in the surface area of fluorescein-stained epithelial defects at both 6 h and 24 h post-wounding (fig. 8A). These results supported that KAMPs can be used to treat infection and inflammation of barrier-compromised sites without the undesirable side effect on wound healing.

**Fig. 8.**
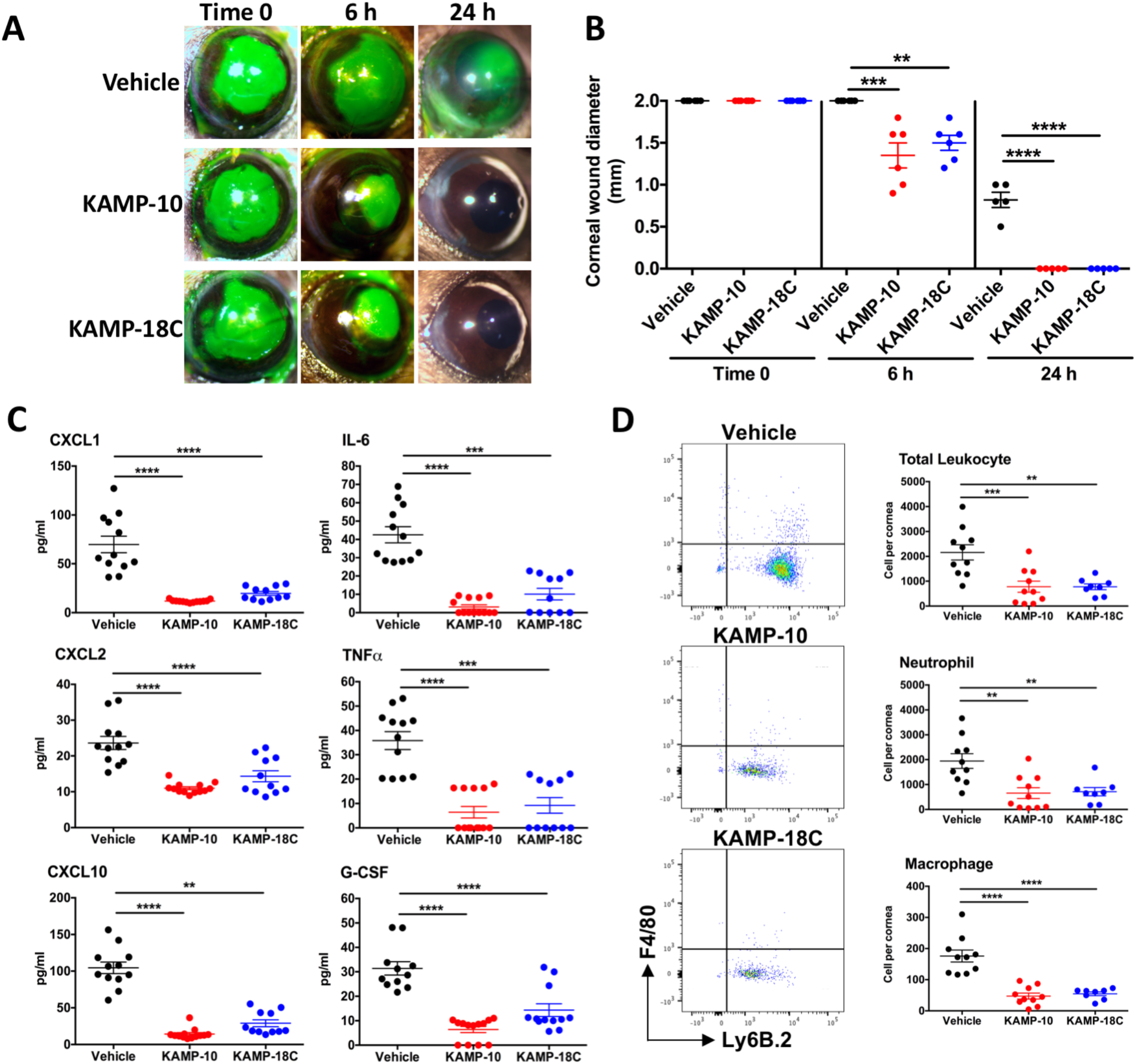
KAMP-10 and KAMP-18C promote corneal epithelial wound healing in a murine model. After removing the corneal epithelium (2 mm diameter), wounds were treated topically with liposome alone (vehicle) or liposome associated KAMP-10 or KAMP-18C (5 μg peptide) three times within 24 h. (**A**) Corneal wounds were stained with fluorescein and photographed. (**B**) Diameter of the wounds at time 0, 6 and 24 h after injury. n=5-6 mice. (**C**) At 6 h post-injury, corneal cytokines were measured by ELISA. n=11-13 mice. (**D**) At 24 h post-injury, single cell suspensions of the corneas were analyzed by flow cytometry to quantify total leukocytes (CD45^+^), neutrophils (CD45^+^Ly6B.2^+^F4/80^-^) and macrophages (CD45^+^F4/80^+^) per cornea. n=8-10 mice. Mean ± SEM are shown. \*\**P* < 0.01, \*\*\**P* < 0.001, \*\*\*\**P* < 0.0001 (ANOVA with Dunnett’s post hoc test except IL-6, TNF*α* and CXCL10, which employed Krustal-Wallis with Dunn’s post hoc test).

## DISCUSSION

Acute inflammatory reactions to bacterial infections are aggressive and must be controlled as early as possible to avoid collateral tissue damage. This is extremely important when treating infection-driven inflammation of delicate organ systems, such as the brain and the eye. To avoid functional loss of the affected organs and achieve best possible clinical outcomes, effective antibiotics and swift alleviation of inflammation are both essential. While corticosteroids are potent first line immunosuppressants, their potential side effects, such as elevated risks of exacerbating infection, warrant careful clinical consideration especially when patients with co-morbidities are involved. Unfortunately, the prudent and timely use of corticosteroids is further complicated by the increasing number of antibiotic-resistant bacteria that are difficult to eradicate. Clearly, effective and optimal treatment of infectious inflammation is facing two distinct but interconnected clinical challenges, both of which could be addressed by new strategies that simultaneously control infection and inflammatory cell infiltration. Using murine sterile and bacterial keratitis models in this study, we have revealed the anti-inflammatory activities of native bactericidal peptides derived from human keratin 6a (KAMPs), and demonstrated their potential as multi-function therapeutic agents that address the unmet needs in the management of acute infectious inflammation through eradication of drug-resistant bacteria, early control of harmful inflammation, and restoration of epithelial integrity.

The first challenge for optimal treatment of infectious inflammation is antibiotic resistance. Each year over two million cases of drug-resistant bacterial infections costing at least 23,000 lives are reported in the United States alone *(51)*. Its negative impact on national and global health has been staggering rapidly and will continue to intensify unless new treatments are made available *(52)*. Furthermore, since drug-resistant bacteria are often more virulent and more inflammatory to innate immune cells, especially after exposure to antibiotics which they have low susceptibility to *(53)*, these infections are far more difficult to control and inevitably delay initiation or increase the risk of adverse effects of adjunctive steroid therapy, leading to substantial burden of morbidity and long-term disability in survivors *(54)*. Antimicrobial peptides (AMPs) are one of the few promising alternatives to conventional antibiotics due to their rapid, multifaceted and broad spectrum bactericidal activities that confer lower propensity for resistance development *(55, 56)*. A number of natural and designed AMPs have been identified or developed over the past three decades and several of them are currently under evaluation in clinical trials for localized infections *(57-59)*. Yet, the main challenges for translating most of the AMP-based drug candidates to the clinical phase of development include their inherent susceptibility to inhibition by in vivo environmental factors, such as physiological salts and proteases, as well as potential cellular toxicity and high cost of production *(25, 60)*.

While technological advances in peptide engineering, nanocarrier formulation and chemical synthesis continue to demonstrate great promise for improving safety, efficacy, stability and manufacturing process of AMPs *(61)*, our study here showed that KAMP-10 and KAMP-18C – the short peptides with native and non-chemically modified sequences from human cytokeratin 6a protein - have no detectable toxic effect on corneal epithelial cells, immune cells and red blood cells; they also remain bactericidal on the ocular surface and in the infected cornea where significant amount of bacterial and host proteases are present *(62, 63)*. It is worth noting that the antibacterial effectiveness of KAMPs was not undermined by the development of antibiotic resistance in the *P. aeruginosa* clinical isolate used in this study. Similar to most peptide-based drugs *(64)*, the major factor that determines therapeutic efficacy of KAMPs is bioavailability at the site of action. With the aid of regular (non-custom-made) phosphatidylcholine/stearylamine/cholesterol liposomes as the topical delivery vehicle, we have been able to forgo invasive subconjunctival injections and yet enhance tissue penetration of KAMPs to the mouse corneal stroma where bacteria and immune cells accumulate during active infection. The proof-of-concept evidence presented here supports further investigations into the range of drug-resistant bacterial species susceptible to KAMPs (alone or in cooperation with current antibiotics), as well as the optimization of physiochemical features of liposomes that enhance efficacy and bioavailability (e.g. tissue penetration for topical use) of KAMPs *(61, 65)*. The prospect of utilizing the antimicrobial characteristic of KAMPs to treat infection is also improved by increasing evidence showing that combined applications of membranolytic AMPs with conventional antibiotics that target intracellular machineries often produce additive or even synergistic bactericidal effects, which in turn afford reduction of drug use and significant potentiation of activity against drug-resistant bacteria *(66)*.

The second challenge for optimal treatment of infectious inflammation is timely control of immune cell influx. While actively participating in innate defense, several AMPs are also known for their roles in immunomodulation since abnormal expression of these peptides are associated with various inflammatory skin and gastrointestinal diseases *(67, 68)*. Typically, these AMPs enhance the recruitment of leukocytes to the site of microbial invasion through their direct chemotactic activity and/or acting on local structural and immune cells to selectively modulate cytokine and chemokine milieu *(69)*. This feature is particularly valuable for clearing drug-resistant bacterial pathogens, as demonstrated by engineered AMPs that reduced pathogen load and mortality in murine peritoneal infection *(70)*. However, the potential risks of collateral tissue damage and resulting functional loss that are associated with amplified inflammatory responses and cellular recruitment remain a major drawback, rendering these AMPs inapplicable to treating infectious inflammation of delicate organs. Indeed, functional outcomes of these diseases are tremendously important, yet often overlooked or insufficiently measured, as most in vivo and ex vivo studies aiming to develop newer and better anti-infectives or anti-inflammatories, including those examining AMPs and their engineered derivatives, have singularly focused on either bacterial killing or amelioration of cytokine production in immune cells for the control of deadly sepsis. In the case of acute infectious inflammation, bifunctional KAMPs can prove particularly useful – not only do KAMPs kill bacteria as demonstrated in the corneal infection model, but the most notable outcome of their concomitant anti-inflammatory activities is the significant reduction and even prevention of immune cell-mediated corneal ulceration and opacification, a leading cause of blindness globally *(71)*. Conceivably, extended studies using animal models of various infectious and non-infectious inflammatory disorders would be necessary to investigate potential indications of KAMPs beyond keratitis for prevention of permanent disabilities.

In this study, we also investigated the anti-inflammatory mechanisms of KAMPs which control bacteria-induced cytokine secretions from neutrophils and macrophages, the cells that make up the first wave of immune infiltrates. KAMPs were found to intervene both the cell surface and the endosomal signaling of TLR2 and TLR4. Instead of direct binding to and neutralization of exogenous bacterial ligands, which is the most common mechanism shared by anti-inflammatory AMPs, KAMPs act as receptor antagonists to simultaneously block multiple cell surface receptors for LPS and LTA. Specifically, KAMPs competitively bind MD-2, the obligatory co-receptor that directly interacts with LPS in the activation of TLR4; TLR2, the most prominent LTA receptor in a heterodimeric complex with TLR1 or TLR6; and CD14, the co-receptor that facilitates the transfer of these two bacterial ligands to their respective cell surface TLR complexes. In addition, KAMPs induce internalization of bacterial ligand-free TLR2 and TLR4, thereby serve to reduce cell surface availability of these receptors before bacterial activation. These findings help explain the suppressive effects of KAMPs on LPS- and LTA-induced NFкB activation. Given that the physical interactions between CD14 and LPS/LTA are required for their induction of TLR2 and TLR4 endocytosis *(41, 72)*, KAMPs competitive binding to CD14 can also explain the inhibition of LPS- and LTA-induced IRF3 activation. Together, the multifaceted mechanisms of KAMPs enable effective prevention or reduction of leukocyte infiltration when these antibacterial peptides are given either prophylactically or therapeutically. Considering that most of the TLR-targeted agents currently in clinical trials are immune stimulatory agonists indicated as vaccine adjuvants or cancer therapies, and that the few immunosuppressive biologics among them are neutralizing antibodies or classic small molecules *(75)*, short bactericidal peptides from human keratin 6a as TLR2/4 antagonists would represent unique drug candidates with great potential to treat both infectious and non-infectious inflammation.

## MATERIALS AND METHODS

### Synthetic peptides

All peptides in HCL salt were synthesized by GenScript USA. Net Peptide content (>70%), peptide purity (>95%) and peptide sequences were confirmed by amino acid analysis, HPLC and mass spectrometry, respectively. Stock solutions (4.8 mg net peptide/ml) were prepared in sterile distilled water and stored at −20°C. Aliquots were limited to one thaw prior to use.

### Bacterial strains

*P. aeruginosa* clinical isolate PA3346 was obtained from a patient diagnosed with bacterial keratitis at the Cole Eye Institute of Cleveland Clinic (Cleveland, OH). *S. aureus* strain 29213 was obtained from ATCC. Amikacin-resistant PA3346 mutant was made by plating 10^9^ CFU on amikacin (50 ug/ml) supplemented-tryptic soy agar (TSA), followed by incubation at 37^°^C overnight. Single colonies of survived bacteria were isolated and grown in amikacin-supplemented tryptic soy broth (TSB) before stocking. To prepare inoculum for infection studies, bacteria were grown in TSB at 37^°^C, 180 rpm for 18 h, then sub-cultured (1:100 dilution in TSB) and grown to log phase (OD650 ∼0.4). Bacteria were washed once and suspended in saline until OD650 ∼0.1 (10^8^ CFU/ml). Bacterial concentrations were determined by serial dilutions and plating on tryptic soy agar.

### Lactate dehydrogenase (LDH) assay

Human telomerase-immortalized corneal epithelial (hTCEpi) cells *(26)* were maintained at 37°C/5% CO2 in regular keratinocyte growth medium KGM-2 (Lonza). Prior to treatment, hTCEpi cells were grown to confluence in 96-well tissue culture-treated plates. To examine the cytotoxic/cytolytic effect of KAMP-10 and KAMP-18C, hTCEpi cells were untreated (baseline control) or treated with various concentrations of KAMPs (25, 50, 100 or 200 μg/ml) for 24 or 48 hours at 37°C/5% CO2. The levels of LDH released into the cell culture medium upon plasma membrane damage were assessed using the colorimetric LDH Cytotoxicity Assay Kit (Thermo Scientific Pierce) according to the manufacturer’s instructions. Cells incubated with lysis buffer served as maximum LDH activity control (100% cytotoxicity).

### Apoptosis Assay

hTCEpi cells were grown to confluency on the Millicell EZ slides with 8 wells (Millipore Sigma). Cells were untreated (baseline control) or treated with DNase (500 U/ml, positive control) or KAMP-10 or KAMP-18C (50, 100 or 200 μg/ml) for 24 h at 37°C/5% CO2. After the treatments, cell nuclei with DNA strand breaks at the early stage of apoptosis were labeled using fluorescein-dUTP in the TUNEL reaction (In Situ Cell Death Detection Kit; Roche) according to the manufacturer’s instructions. Cell nuclei were also stained with DAPI before viewing under a phase contrast/fluorescence microscope (Zeiss Axio Observer).

### Hemolytic Assay

Single donor red blood cells (RBCs) were purchased from Innovative Research Inc. 5×10^6^ cells in 50 μl of PBS were placed in 96-well plates, topped with 50 μl of PBS (baseline), or PBS containing 0.1% Triton X-100 (complete hemolysis as positive control), or KAMP-10 or KAMP-18C at 25, 50, 100, 200 or 400 μg/ml. Samples were incubated for 3 hours at 37°C and agitated every 10 min. Immediately after incubation, supernatants (100 μl) were collected and the absorbance of liberated hemoglobin was measured at 540 nm. Percent of hemolysis = A1/A2 × 100, where A1 was absorbance of sample and A2 was absorbance of positive control.

### Murine models of corneal inflammation, infection or wound healing

All procedures were conducted in compliance with the Public Health Service Policy on Humane Care and Use of Laboratory Animals, and approved by the Institutional Animal Care and Use Committee of the Cleveland Clinic. Both female and male C57BL/6J mice (8-10 weeks old) were anesthetized by intraperitoneal injection (50 μl per 25 g body weight) of ketamine (25 mg/ml) and dexmedetomidine (0.2 mg/ml) cocktail. To compromise the epithelial barrier, three parallel abrasions were made on the mouse corneas using a 26g needle. In the sterile inflammation model, purified *P. aeruginosa* LPS or *S. aureus* LTA (20 μg in 2 μl PBS; Sigma-Aldrich), or PBS only (scratch control), was topically applied to the scarified corneas. LPS- or LTA-simulated mice were either mock-treated or topically applied once (5 μl) with KAMP-10 or KAMP-18C 30 min before (100 μg/ml) or after (200 μg/ml) stimulation. In the bacterial infection model, live amikacin-resistant *P. aeruginosa* PA3346 (10^5^ CFU in 5 μl saline) or *S. aureus* (10^6^ CFU in 5 μl saline) was inoculated on the scarified corneas. For infection prevention, KAMP-10 or KAMP-18C (200 ng) was topically applied once at the time of inoculation. To prepare treatment for ongoing infection, a liposome kit with lyophilized lipid powder containing L-*α*-phosphatidylcholine (63 μmoles), stearylamine (18 μmoles) and cholesterol (9 μmoles) (Sigma Aldrich) was reconstituted with 300 μl of saline, following by gentle mixing with 700 μl of saline (vehicle control), or KAMP-10 or KAMP-18C (1.5 mg/ml saline). The resulting mixture (5 μg peptide in 5 μl) was then topically applied to infected corneas three times daily for 3 days beginning at 24 h post-infection. Infected corneas were examined daily under a stereomicroscope equipped with digital camera to monitor disease progression. Area of opacity, density of central opacity, density of peripheral opacity and surface regularity were blind graded with 0-4 points each to determine total disease scores (0-16 points). In the corneal wound healing model, the epithelium (2 mm diameter) at the central cornea was gently removed using a Beaver6400 mini-blade (Beaver Visitec). Fluorescein sodium 0.4% was used to stain compromised barrier immediately, 6 h and 24 h after injury. At experimental endpoint, whole corneas were dissected with a trephine and a surgical microscissor and homogenized in sterile PBS (150 μl each) using Precellys CK14 lysing kit and tissue homogenizer (Bertin Instruments). After centrifugation to remove debris, bacterial loads were quantified by serial dilution and plating on tryptic soy agar, and cytokine levels were determined by ELISA (DuoSet, R&D Systems) in the half area 96-well microplate format (Corning Costar). For flow cytometric analysis of immune cells, mouse corneas were incubated in type I collagenase (Sigma Aldrich) at 82 U/cornea for 2 hours at 37^°^C. Cell suspensions were passed through 30 μm filter once to remove any undigested tissue.

### Flow cytometric analysis of neutrophils and macrophages

C57BL/6J mice (Jackson Laboratory) were euthanized immediately prior to the collection of neutrophils and macrophages. For bone marrow neutrophils, total bone marrow cells were collected from tibias and femurs, then resuspended in 1 ml of sterile RBC lysis buffer (Sigma-Aldrich) and placed on ice for 10 minutes. Remaining cells were washed twice with PBS, then processed with EasySep Mouse Neutrophil Enrichment Kit (StemCell Technologies) to isolate neutrophils by negative selection. For thioglycollate-elicited peritoneal cells, C57BL/6J mice were intraperitoneally injected with 1 ml of 3% Brewer thioglycollate medium (Sigma-Aldrich). Elicited peritoneal neutrophils and macrophages were collected 1 day and 3 days after the injection, respectively, by injecting 10 ml of sterile PBS into the peritoneal cavity, followed by gently massaging the peritoneum, then collecting with a 18g needle/syringe. Cells in PBS were pelleted and resuspended in RBC lysis buffer for 10 min. Typically, primary neutrophils and macrophages were resuspended in DMEM and RPMI respectively (both containing 2% FBS), and placed in 96-well plate (1×10^5^ cells/well). Macrophages were additionally cultured at 37°C for 6 h to select adherent cells for subsequent treatments. Bone marrow neutrophils and resident peritoneal macrophages were untreated, or treated with LPS or LTA (500 ng/ml) in the presence or absence of KAMP-10 or KAMP-18C (50, 100 or 200 μg/ml) for 24 h at 37°C/5% CO2. KAMPs were added 30 min either before or after bacterial ligand stimulation. Cytokines in cell culture supernatants were measured by ELISA (DuoSet; R&D) or mouse 31-plex cytokine/chemokine arrays (Eve Technologies). Thioglycollate-elicited neutrophils and macrophages were treated with 500 U/ml of IFN-γ for 18 h, washed, then either unstimulated or stimulated with 500 ng/ml LPS or LTA, or 200 ug/ml of KAMP-10, KAMP-18C or SC-10, alone or in combination, for 1 hour. For flow cytometry, cells were washed and incubated on ice for 10 min in 100 μl ice cold FACS buffer (PBS with 1% FBS) containing 2 μg Fc blocker (anti-mouse CD16/CD32 antibody; eBioscience). Antibody staining was typically conducted on ice for 1 h with rat monoclonal FITC or PE anti-mouse Ly6B.2 (clone 7/4; Abcam), FITC or PE/Cy5 anti-mouse F4/80 (clone BM8; Biolegend), APC/Cy7 anti-mouse CD45 (clone 30-F11; Biolegend), APC or PE anti-mouse TLR4 antibody (clone SA15-21; Biolegend), recombinant PE anti-human/mouse TLR2 (clone QA16A01; Biolegend), FITC anti-mouse CD14 (clone Sa14-2; Biolegend), rabbit monoclonal Alexa 647 anti-phospho-p65(Ser536) (clone 93H1; Cell Signaling) or anti-phospho-IRF3(Ser 396) (clone D6O1M; Cell Signaling) antibodies or the corresponding isotype controls. All cells were also stained with Zombie Violet (Biolegend) to gate out dead cells. Stained cells were washed twice with FACS buffer and resuspended in 0.5% PFA for detection with a BD LSRII flow cytometer. Cells were first gated using forward scatter (FSC) and side scatter (SSC), then gated on viability, CD45+ leukocyte population, and cell type markers (Ly6B.2 and F4/80). Unstained samples, isotype controls and florescence minus one were used to confirm the gating strategy. Flow cytometric analysis was performed using BD FlowJo software.

### Flow cytometric analysis of TLR-conjugated beads

Carboxylate-modified polystyrene latex (CML) beads of 7 μm diameter (ThermoFisher) were conjugated with recombinant TLR as previously described *(76)*. Briefly, 18×10^6^ beads in 50 μl MES coupling buffer (50 mM MES (pH 6.0) with 1 mM EDTA) were incubated with 20 μl of EDAC (50 mg/ml in MES coupling buffer) for 15 min at 25^°^C to activate the carboxyl groups on the beads. After extensive washing in PBS, activated beads were resuspended in 200 μl PBS. One quarter of the bead suspension was mixed with 50 μl of PBS containing 25 μg of recombinant human TLR4/MD-2 complex or TLR2 (R&D) for 3 hours at 25^°^C. Following conjugation, beads were washed with PBS and stored in 100 μl of QBS buffer (PBS containing 1% BSA and 0.02% sodium azide). To assess KAMPs binding to TLR4/MD-2, TLR4/MD-2 complex-conjugated beads (5,000 in FACS buffer) were preincubated with KAMP-10 (48 μg/ml), KAMP-18C (96 μg/ml) or SC-10 (48 μg/ml) in the presence or absence of recombinant human MD-2 (R&D) (1 μg/ml) for 1 h on ice. Beads were then washed twice with PBS before incubation with FITC-E. coli O55:B5 LPS (1 μg/ml) (Sigma-Aldrich) for 1 h on ice. Beads were washed with PBS again, and TLR4 on the beads were stained with mouse monoclonal APC anti-human TLR4 antibody (clone HTA125; ThermoFisher) for 1 h on ice prior to flow cytometric analysis. Similarly, to assess KAMPs binding to TLR2, TLR2-conjugated beads preincubated with individual KAMP at the aforementioned concentrations were washed with PBS, then incubated with *S. aureus* LTA (1 μg/ml) (Sigma-Aldrich) and recombinant human CD14 (R&D) (0.1 μM) for 1 h. To assess the effect of KAMPs on CD14-LTA interaction, KAMPs (0.1 μM equivalent to 78 ng/ml for KAMP-10 and SC-10; 160 ng/ml for KAMP-18C) were preincubated with equal molar of CD14 (0.1 μM) for 1 h on ice before incubation with TLR2-conjugated beads and LTA for 1 h. To detect LTA and TLR2 by flow cytometry, mouse monoclonal FITC anti-LTA antibody (clone 55; Novus Biologicals) and mouse recombinant PE anti-human/mouse TLR2 (clone QA16A01; Biolegend) were used.

### Molecular modeling

The HPEPDOCK server *(77)* was employed to perform blind global docking of KAMP-10 and KAMP-18C sequences to crystal structures of mouse TLR4/MD-2 complex (PDB ID: 5IJB), human MD-2 (PDB ID: 2E56), mouse TLR2 (PDB ID: 3A79), and human CD14 (PDB ID: 4GLP) through a hierarchical flexible peptide docking approach and global sampling of binding orientations.

### Statistical analysis

GraphPad Prism was used to perform statistical tests. Data normality was determined by D’agostino-Pearson omnibus test. For normally distributed data, unpaired two-tailed Student’s t-test and one-way ANOVA with post hoc Dunnett’s test were used for single and multiple intergroup comparisons, respectively. For multiple intergroup comparisons of non-normally distributed data, Krustal Wallis with post hoc Dunn’s test was used. A significance level of *P* < 0.05 was considered statistically significant.

## SUPPLEMENTARY MATERIALS

**Fig. S1. Dose-dependent suppression of LPS- and LTA-induced CXCL10, TNF*α*, and G-CSF secretion from primary mouse neutrophils and macrophages by KAMP-10 and KAMP-18C**.

**Fig. S2. Multiplex analysis of LPS- and LTA-induced cytokine production from primary mouse neutrophils treated with a sub-effective dose of KAMP-10 or KAMP-18C**.

**Fig. S3. Multiplex analysis of LPS- and LTA-induced cytokine production from primary mouse macrophages treated with a sub-effective dose of KAMP-10 or KAMP-18C**.

**Fig. S4. Immunofluorescence microscopy of neutrophil infiltration in LPS-stimulated mouse corneas either untreated or treated with KAMP-10**.

## Author contributions

YS, JC and KB performed the experiments. CT and YS designed the experiments, analyzed and interpreted the data, and wrote the manuscript. CT conceived and supervised the project.

