## Supplemental Figures for "Simultaneous Control of Infection and Inflammation by Keratin-Derived Antibacterial Peptides (KAMPs) Targeting TLRs and Co-Receptors"

### Supplemental Figure 1

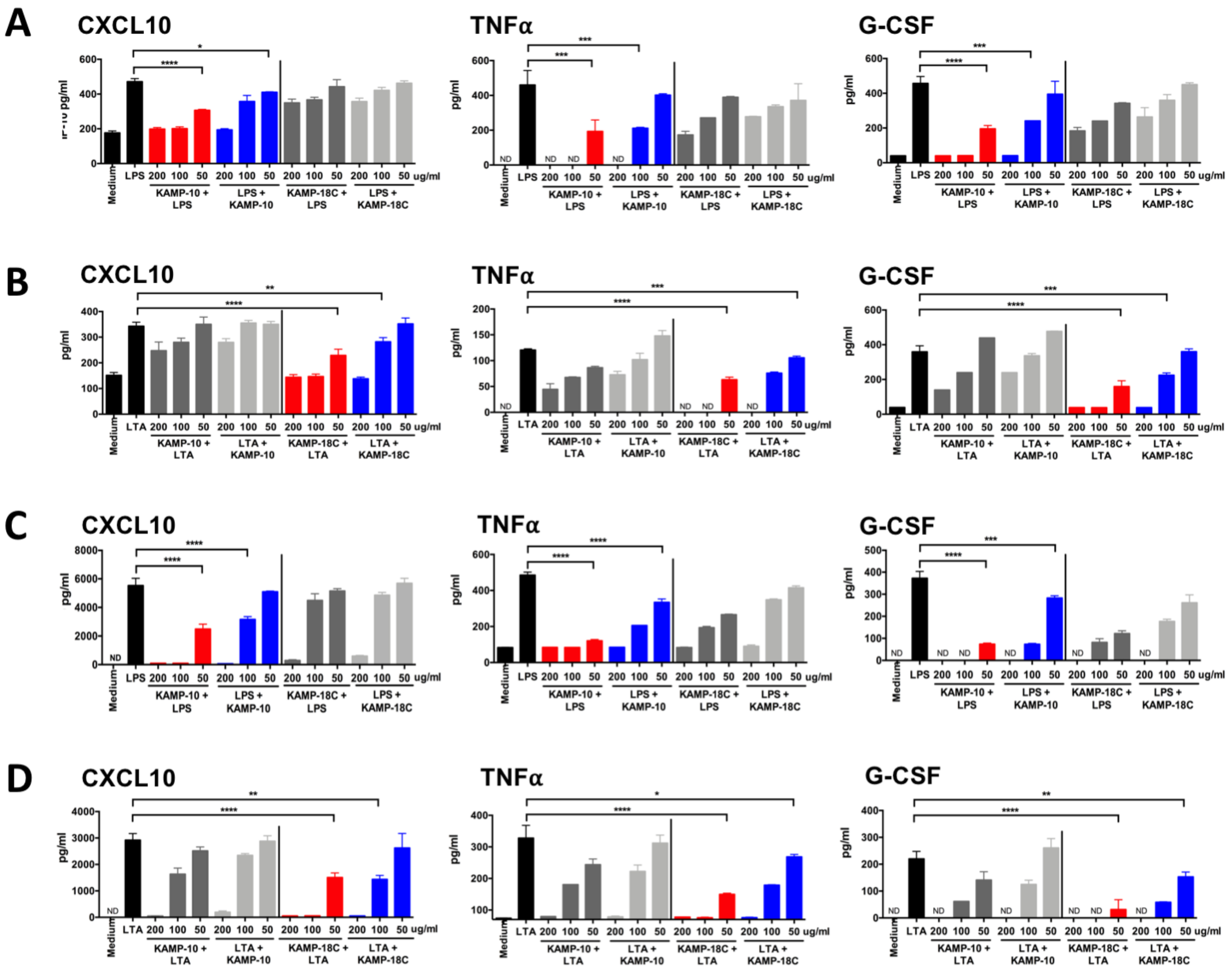

**Fig. S1. Dose-dependent suppression of LPS- and LTA-induced CXCL10, TNFα, and G-CSF secretion from primary mouse neutrophils and macrophages by KAMP-10 and KAMP-18C.** (**A-B**) Enriched murine bone marrow neutrophils and (**C-D**) resident peritoneal macrophages were stimulated with LPS (A, C) or LTA (B, D) (500 ng/ml) in the absence or presence of KAMP-10 or KAMP-18C at the indicated concentrations for 24 hours. The peptides were applied either 30 minutes before (red and dark grey) or 30 minutes after (blue and light grey) the stimulation. Cells mock-treated with medium served as baseline controls. CXCL10, TNFα, G-CSF levels in culture supernatants were measured by ELISA. Mean of three replicates ± SD are shown. ND=non-detected. *$P < 0.05$, **$P < 0.01$, ***$P < 0.001$, ****$P < 0.0001$ (ANOVA with Dunnett's post hoc test for multiple comparisons). The lowest concentration of KAMP yielding statistical significance are shown.

#### Supplemental Figure 2

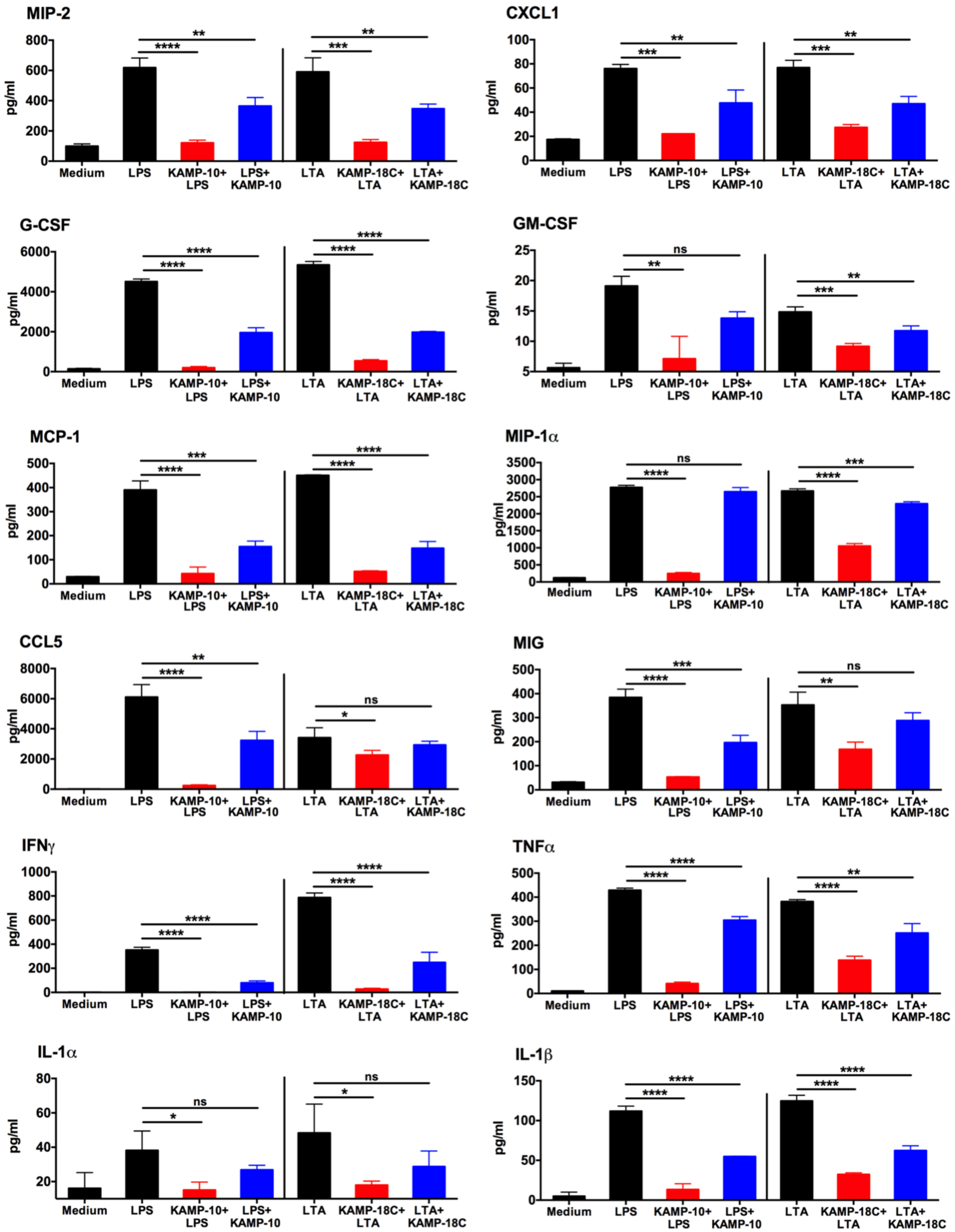

**Fig. S2. Multiplex analysis of LPS- or LTA-induced cytokine production from primary mouse neutrophils treated with a sub-effective dose of KAMP-10 or KAMP-18C.** Enriched murine bone marrow neutrophils were stimulated with LPS (500 ng/ml) in the absence or presence of KAMP-10, or LTA (500 ng/ml) in the absence or presence of KAMP-18C (50 μg/ml). KAMPs were added to culture media either 30 min before (KAMP+bacterial ligand) or 30 min after stimulation (bacterial ligand+KAMP). Cells mock-treated with medium served as baseline controls. Culture supernatants were collected 24 hours after stimulation and analyzed by fluorescent bead-based multiplex ELISA. Mean (n=3 replicates) ± SD are shown. $^*P < 0.05$, $^{**}P < 0.01$, $^{***}P < 0.001$, $^{****}P < 0.0001$, ns = non-significant (ANOVA with Dunnett's post hoc test).

#### Supplemental Figure 3

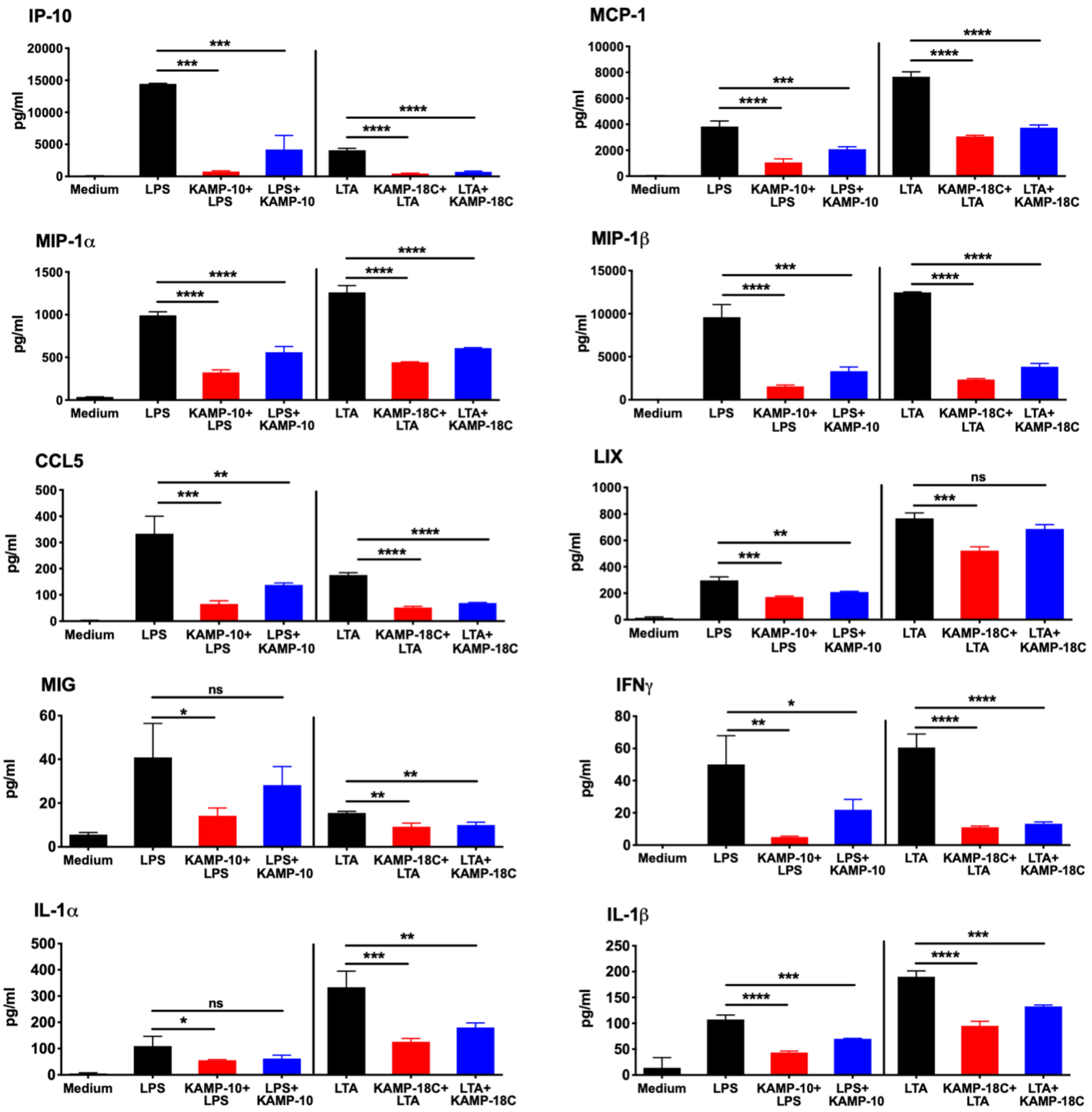

**Fig. S3. Multiplex analysis of LPS- or LTA-induced cytokine production from primary mouse macrophages treated with a sub-effective dose of KAMP-10 or KAMP-18C.** Enriched murine resident peritoneal macrophages were stimulated with LPS (500 ng/ml) in the absence or presence of KAMP-10 (50 μg/ml), or LTA (500 ng/ml) in the absence or presence of KAMP-18C (50 μg/ml). KAMPs were added to culture media either 30 min before (KAMP+bacterial ligand) or 30 min after stimulation (bacterial ligand+KAMP). Cells mock-treated with medium served as baseline controls. Culture supernatants were collected 24 hours after stimulation and analyzed by fluorescent bead-based multiplex ELISA. ELISA. Mean (n=3 replicates) ± SD are shown. *$P < 0.05$, **$P < 0.01$, ***$P < 0.001$, ****$P < 0.0001$, ns = non-significant (ANOVA with Dunnett's post hoc test).

#### Supplemental Figure 4

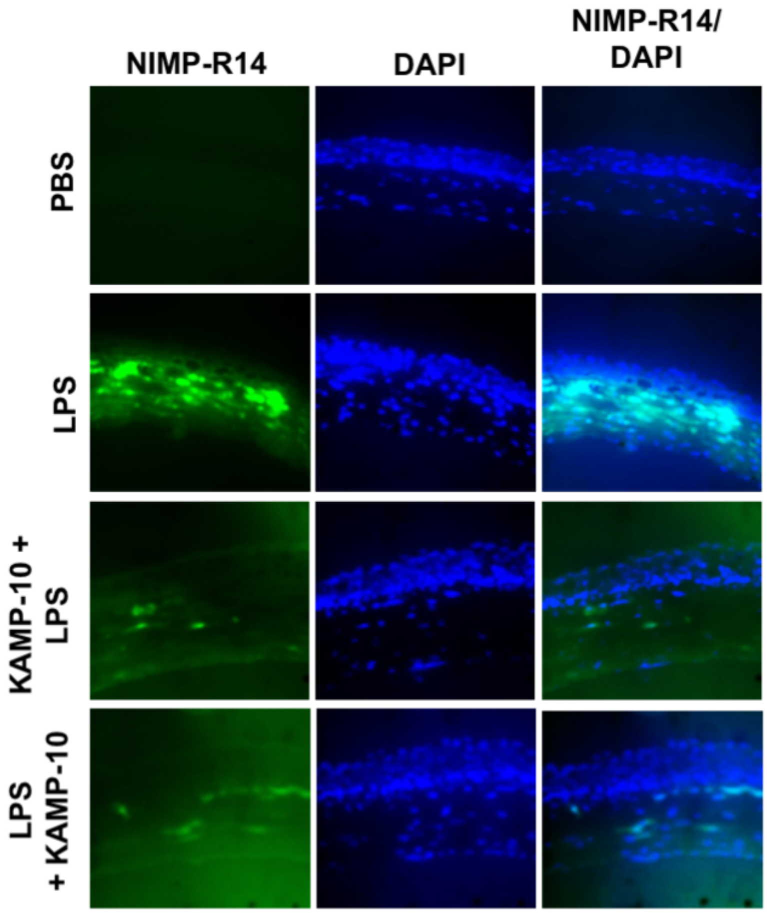

**Fig. S4. Immunofluorescence microscopy of neutrophil infiltration in LPS-stimulated mouse corneas either untreated or treated with KAMP-10.** In a mouse model of sterile corneal inflammation, purified *P. aeruginosa* LPS (20 μg in 2 μl PBS) or PBS only (2 μl) was topically applied to scarified mouse corneas. LPS-inoculated mouse eyes were untreated, or treated with one topical application (5 μl) of KAMP-10 either 30 minutes before (100 μg/ml KAMP-10 + LPS) or 30 minutes after LPS inoculation (LPS + 200 μg/ml KAMP-10). After 24 hours, mice were euthanized and whole eyes were frozen in optimal cutting temperature (OCT) embedding medium before cross-sectioning to 7 μm thickness with a cryostat. Sections were fixed in 4% formaldehyde for 20 minutes, and neutrophils (green) were stained for 2 h by rat anti-mouse Ly-6G/-6C antibody NIMP-R14 (Abcam; 1:100 dilution in TBS) and 45 min by FITC-conjugated rabbit anti-rat antibody (Vector Laboratories; 1:100 dilution in TBS) at room temperature. Cell nuclei (blue) were counterstained and mounted with Vectashield antifade medium containing DAPI (Vector Laboratories), then viewed under a fluorescence microscope (Zeiss Axio Observer).
